# A subcellular biochemical model for T6SS dynamics reveals winning competitive strategies

**DOI:** 10.1101/2021.07.17.452664

**Authors:** Yuexia Luna Lin, Stephanie N. Smith, Eva Kanso, Alecia N. Septer, Chris H. Rycroft

## Abstract

The Type VI secretion system (T6SS) is a broadly distributed interbacterial weapon that can be used to eliminate competing bacterial populations. Although unarmed target populations are typically used to study T6SS function, bacteria most likely encounter other T6SS-armed competitors in nature. The outcome of such battles is not well understood, neither is the connection between the outcomes with the subcellular details of the T6SS. Here, we incorporated new biological data derived from natural competitors of *Vibrio fischeri* light organ symbionts to build a biochemical model for T6SS function at the single cell level. The model accounts for activation of structure formation, structure assembly, and deployment. By developing an integrated agent-based model (IABM) that incorporates strain-specific T6SS parameters, we replicated outcomes of biological competitions, validating our approach. We used the IABM to isolate and manipulate strain-specific physiological differences between competitors, in a way that is not possible using biological samples, to identify winning strategies for T6SS-armed populations. We found that a tipping point exists where the cost of building more T6SS weapons outweighs their protective ability. Furthermore, we found that competitions between a T6SS-armed population and a unarmed target had different outcomes dependent on the geometry of the battlefield: target cells survived at the edges of a range expansion scenario where unlimited territory could be claimed, while competitions within a confined space, much like the light organ crypts where natural *V. fischeri* compete, resulted in the rapid elimination of the unarmed competitor.

---

All life forms compete with one another in what Charles Darwin referred to as “the struggle for life” [1]. Indeed, fierce battles for limited space and resources can be observed across biological complexity, from single cell organisms to humans. These battles often determine which population will proliferate, and which will be excluded. Therefore, organisms at all size scales have evolved diverse strategies to wage war on their rivals and increase the probability of their own success.

Microbial genomes encode an incredible arsenal of interbacterial weaponry [2], and some, e.g., the broadly distributed type VI secretion system (T6SS), have been shown to be useful in competing for colonization sites within a host niche [3, 4, 5, 6, 7, 8, 9]. T6SSs resemble a molecular syringe and are thought to have evolved from bacteriophage contractile tails [10, 11]. T6SS-containing cells build a sheath-and-tube structure that extends the width of a cell and is anchored into the cell wall and membrane [12]. When the sheath is contracted, the inner tube and toxins are propelled into the neighboring target cell, resulting in death and lysis of the competitor [13, 14, 15, 16].

Although genome sequencing allows us to identify the bacterial strains that harbor T6SS interbacterial weapons [17], we are unable to predict which microbial population will dominate in an ecologically relevant battle based on genetic data alone, because the details of when and how such weapons are deployed are lacking. Thus, mathematical and numerical models hold the key to understanding the dynamics of T6SS weapons and predicting its impact on the microbial populations at large, because they allow us to manipulate the system with a larger degree of freedom.

Among the various models for microbial activities, approaches based on ordinary and partial differential equations have been fruitful in predicting T6SS-dependent competition outcomes [18]. In parallel, the agent-based models (ABMs), which have long been used to study population dynamics and spatial organizations in diverse contexts [19, 20, 21, 22], have also been applied to model T6SS-dependent competitions between bacterial populations [23, 24, 15, 25] and yielded predictions including T6SS effectiveness depends on the speed of target cell lysis induced by the T6SS toxins [15]. Existing T6SS ABMs have largely focused on the “rules of engagement” among cells, e.g. the frequency of firing, the probability of hitting a target, and the time it takes for the attacked to disintegrate. Here, we expand the use of ABMs by incorporating a subcellular biochemical model of T6SS that accounts for how T6SS activity and the number of weapons in the arsenal are regulated within a cell. We identify new T6SS-related parameters and quantify them using biological experiments with *Vibrio fischeri* light organ isolates, which are natural competitors of the squid light organ niche [7, 26, 27]. Using this combined model, we are able to replicate the impact of T6SS-dependent killing on population dynamics and the spatial distribution of competing strains observed in experiments. Furthermore, our model offers novel hypotheses on how factors such as strain-specific physiological differences and spatial geometry, impact competitive outcomes in ecologically relevant settings. Although engineering lab mutants have become standard research practice, there are yet limitations in practical feasibility and controllability when probing a biological system. Our model thus provides a viable alternative approach to generate and test hypotheses about T6SS activity that would be difficult or impossible to address experimentally.

## Results

### Competition outcomes vary due to intraspecific variations in T6SS killing dynamics

The ability of lethal strains of *V. fischeri*, which encode a strain-specific T6SS genomic island (T6SS2), to produce T6SS structures and kill target cells is dependent on environmental stimuli [28]. In liquid culture, *V. fischeri* T6SS is functionally inactive, while the exposure to a high viscosity medium such as hydrogel or an agar surface causes the cells to activate T6SS protein expression and structure formation [28]. We hypothesized that the response time to surface activation may differ among lethal strains of *V. fischeri*, and that these variations may affect competition outcomes between two lethal strains. To begin testing this hypothesis, we coincubated the T6SS2-encoding *V. fischeri* strains ES401 and FQ-A002 [7] on LBS agar plates following two different preparations: clonal cultures of each strain were first incubated for 6 h either in (1) LBS liquid medium where T6SS2 activity is low, or (2) on LBS agar plates where T6SS2 activity is increased. Treatments where strains were incubated in liquid are referred to as “unprimed” because both strains come from a “T6SS off” condition and must activate their T6SS at the start of their coincubation on agar surfaces, whereas strains incubated on agar are referred to as “primed” because these strains have fully activated T6SSs when they begin their coincubation on agar surfaces. We predicted that if response time to surface activation is different between these two strains, then one strain will dominate in only the unprimed condition, where it has the advantage of more quickly activating its T6SS to begin killing its competitor, which has a delayed activation time.

When unprimed wildtype ES401 and FQ-A002 were coincubated on LBS agar plates, microscopy images and colony forming unit (CFU) counts revealed that ES401 out-competed FQ-A002 at the population level and only small microcolonies of FQ-A002 remained after 24 h of coincubation (Fig. 1AD). However, when primed wildtype ES401 and FQ-A002 were coincubated, the two strains coexisted with comparable CFU counts and formed distinct, spatially separated microcolonies (Fig. 1BD). To determine whether this effect was dependent on T6SS2 activity, we disrupted the *vasA_2* gene in both ES401 and FQ-A002, which encodes a baseplate protein in the T6SS2 complex and is required for T6SS-dependent killing [7, 28]. These *vasA* mutant strains were then coincubated in both primed and unprimed conditions. We observed that regardless of coincubation conditions, both *vasA*^−^ strains coexisted and were well-mixed, with no spatial separation between strains (Fig. 1CD; SI Appendix Fig. S1A). These findings, which reveal that one lethal strain (ES401) can dominate over another (FQ-A002) when they are forced to activate their T6SS at the onset of competition, support our hypothesis that surface activation timing may impact competitive outcomes and warrant further investigation.

**Figure 1:**
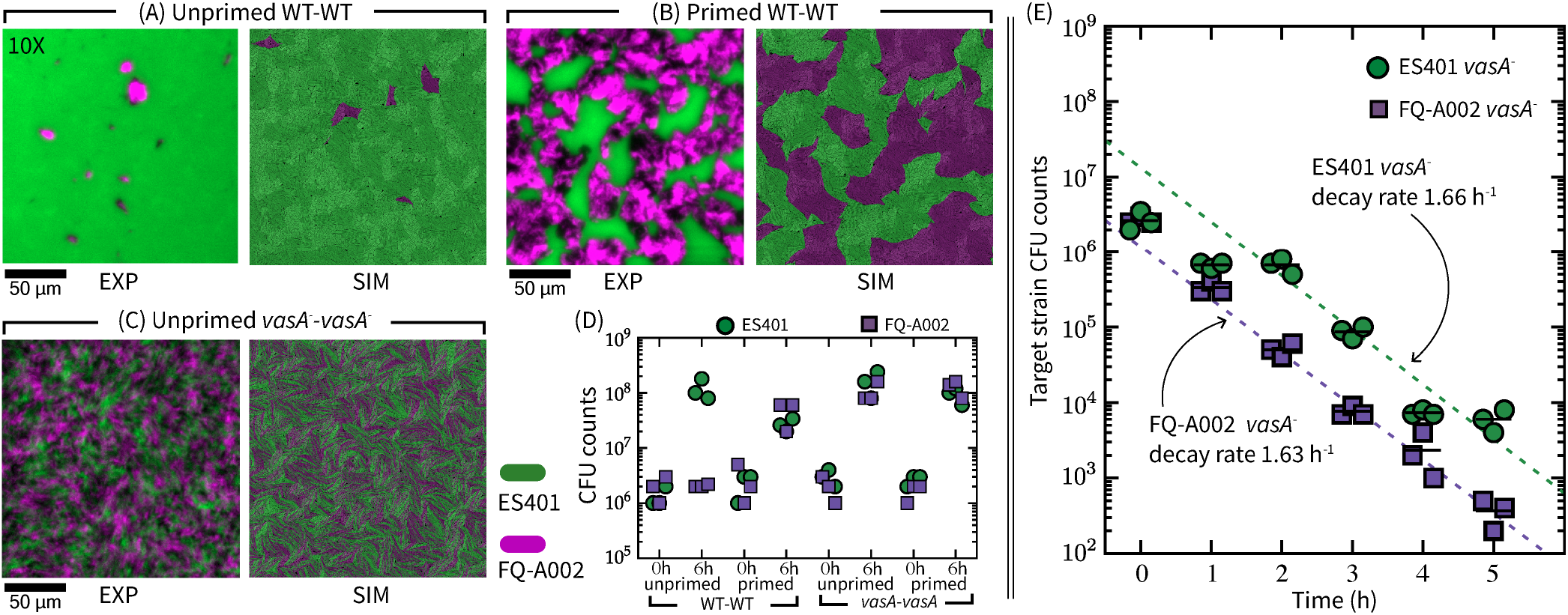
Competition outcomes vary due to intraspecific variations in T6SS killing dynamics. Fluorescence microscopy images of ES401 and FQ-A002 strains at 24 h following coincubation on LBS agar, compared side by side with representative simulation images from our ABM with internal T6SS dynamics (details in text). Microscopy images were taken at 10X; scale bars are 50 μm for all microscopy and simulation images. Coincubations included wildtype vs. wildtype under unprimed treatment (A) and primed treatment (B), and a *vasA*^−^ vs. *vasA*^−^ pair under unprimed treatment (C). The *vasA*^−^ vs. *vasA*^−^ coincubation under primed treatment gives similar outcome to (C) (SI Appendix Fig. S1A). The ES401 strain harbored the GFP-encoding plasmid pVSV102 (green) and the FQ-A002 strain harbored the dsRed-encoding plasmid pVSV208 (magenta). (D) Total colony forming unit (CFU) counts of each strain in each coincubation at 0 h and 6 h following coincubation on LBS agar. (A) - (D) All experiments were performed three times (*n* = 3), all simulations repeated 5 times, representative data are shown. (E) Total target strain CFU counts taken each hour over the course of a 5 h unprimed coincubation between either wildtype FQ-A002 and an ES401 *vasA*^−^ mutant, or wildtype ES401 and an FQ-A002 *vasA*^−^ mutant. Experiment was performed three times (n = 3), all data are shown. Additional simulation data see SI Appendix Fig. S1, SI movies S1, S2.

Based on the above findings, we reasoned that ES401 may activate its T6SS, and begin killing target cells, before FQ-A002, thus providing it an advantage in the competition. To test whether strains ES401 and FQ-A002 have different killing dynamics under unprimed conditions, we directly quantified their ability to eliminate a target population over time. Specifically, we hypothesized that a target population could survive longer and in larger numbers when coincubated with a slower activating lethal strain. To test this hypothesis, we competed either (1) wildtype FQ-A002 vs. ES401 *vasA*^−^, or (2) wildtype ES401 vs. FQ-A002 *vasA*^−^ strains by spotting unprimed mixtures of each treatment on LBS agar pads and obtaining CFU counts of the lethal (wildtype) and the target (*vasA*^−^) strain every hour during a 5 h coincubation period. When the target strain CFUs were plotted over time for each coincubation, we observed that the target populations in both treatments declined exponentially at approximately the same rate, i.e. ~*e*^−1.6*t*^, where *t* is time after spotting (Fig. 1E), suggesting both of the wildtype ES401 and FQ-A002 lethal strains kill at a comparable rate. However, we observed that there was an approximately 10-fold drop in target CFUs between 1 h–2 h in coincubations between wildtype ES401 vs. FQ-A002 *vasA^−^* target, whereas the ES401 *vasA*^−^ target CFUs were maintained between 1 h–2 h in coincubation with the wildtype FQ-A002 (Fig. 1E), suggesting that ES401 begins killing target before FQ-A002. When the CFUs of wildtype ES401 and wildtype FQ-A002 were plotted, we found that both strains exhibit similar growth rates under these conditions (SI Appendix Fig. S2); thus, a difference in growth rate does not account for the differences in target decline between treatments.

Taken together, these results suggest that wildtype ES401 is more effective in eliminating nonlethal targets, when compared to wildtype FQ-A002. Combined with our previous observation that wildtype ES401 outcompetes wildtype FQ-A002 in an unprimed competition, these results support our hypothesis that variations in competitive outcomes are driven by a strain-specific T6SS activation response to surfaces. To investigate this surface response in *V. fischeri*, we performed further experiments to quantify the percentage of cells in a population with T6SSs and the number of structures per cell over time.

### Quantifying T6SS activation dynamics reveals strain-specific differences

In a T6SS complex, VipAB/TssBC multimers comprise the outer component of the sheath structure; thus, VipA has been used in multiple systems as a target for visualization of sheath dynamics through the use of a fluorescently-tagged VipA/TssB fusion constructs [14, 29, 30], including an IPTG-inducible VipA-GFP expression vector in *V. fischeri* [7, 31]. To visualize T6SS activation dynamics, we grew overnight cultures of wildtype ES401 or FQ-A002 strains harboring the VipA-GFP expression vector to an OD_600_ ~ 1.5 and spotted them onto an agarose pad supplemented with 0.5 mM IPTG immediately prior to imaging. FITC images of either ES401 or FQ-A002 expressing VipA-GFP were taken for these analyses (Fig. 2A). We categorized a cell as T6SS activated when we observed at least one sheath within the cell. To quantify the rate of T6SS activation for each strain, we measured the proportion of activated cells in the FITC images taken at regular intervals between 0.5 h and 6 h after initial spotting (Fig. 2B). These measurements revealed that the proportion of activated cells in both ES401 and FQ-A002 remained stable at low levels for approximately 1 h after plating. Directly from the liquid cultures, the activated cells are approximately 10% and 5%for ES401 and FQ-A002, respectively. After this initial waiting period, we observed the activated proportion increased over time in both ES401 and FQ-A002 but the rate of increase was approximately twice as high in ES401 compared to FQ-A002, suggesting ES401 activates T6SS structure assembly more quickly than FQ-A002.

**Figure 2:**
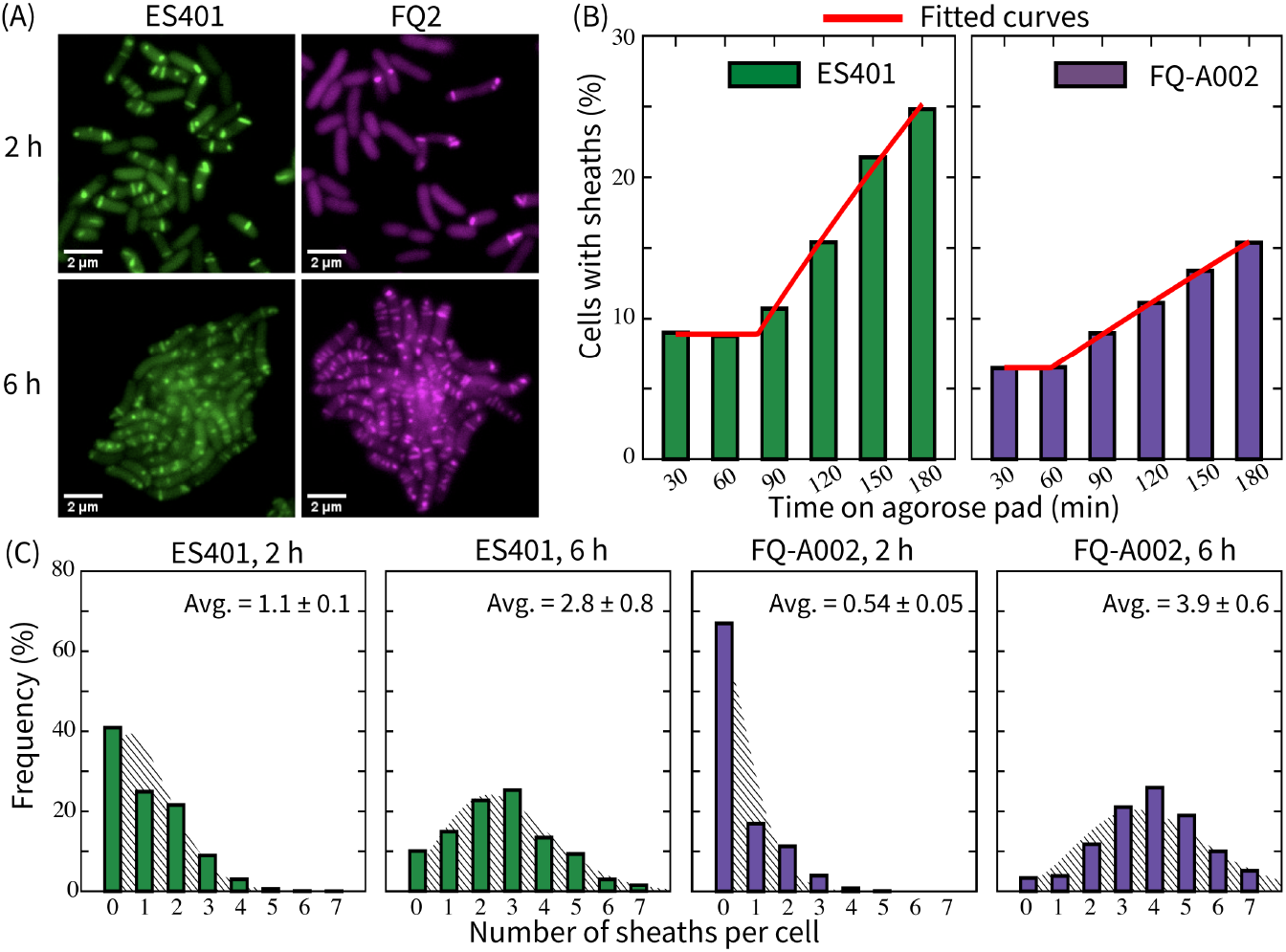
*V. fischeri* exhibits strain-specific T6SS dynamics over time during surface activation. (A) Representative FITC images of ES401 or FQ-A002 cells harboring an IPTG-inducible VipA-GFP expression vector after incubation on an agarose pad for either 2 h or 6 h. Lookup tables (LUTs) were assigned in post-processing to differentiate between *V. fischeri* cells: ES401 was assigned a green LUT and FQ-A002 was assigned a magenta LUT. (B) Percentage of ES041 or FQ-A002 VipA-GFP-expressing cells that contain at least one sheath at 30 min intervals for 3 h. A minimum of 680 total cells were analyzed for each treatment across five fields of view for two biological replicates. All combined data is shown. Parameters in Eq. (2) are estimated to be (*p*_0_, *τ*_+_, λ_+_) = (10%, 1.34 h, 0.118 h^−1^) for ES401, and (*p*_0_,*τ*_+_, λ_+_) = (5%, 1 h, 0.05 h^−1^) for FQ-A002. (C) Distribution of the number of VipA-GFP sheaths per cell in either ES401 or FQ-A002 following either a 2 h or a 6 h incubation on an agarose pad. These data are overlaid with a Poisson distribution with the corresponding mean. > 2700 cells are analyzed for sheath distributions at 2 h, > 260 cells are analyzed at 6 h. All agarose pads were made by supplementing liquid LBS with 2% agarose and 0.5 mM IPTG (isopropyl-*β*-D-thiogalactopyranoside). (A) and (C) show data from the same experiments, which are performed two times and all combined data are shown.

We reasoned that the number of weapons in a strain’s T6SS arsenal may also impact competition outcomes in either positive or negative ways, as more weapons might allow a strain to kill faster, but this advantage could come at an energetic cost. Therefore, we next quantified the average number of sheaths per cell over time after *V. fischeri* cells are spotted on agar surfaces. We incubated ES401 and FQ-A002 clonally on an agarose surface as described above. For each strain, we counted the number of sheaths per cell for a given population in the FITC images at 2 h and 6 h and plotted the frequency of having 0–7 sheaths at each time point (Fig. 2C). For both strains, the average number of sheaths per cell increased between 2 h and 6 h on surfaces. However, this value was different for each strain: ES401 went from an average of 1.1 sheaths per cell at 2 h to 2.8 sheaths per cell at 6 h, while FQ-A002 went from 0.5 at 2 h to 3.9 sheaths per cell at 6 h. We also found that the experimentally measured distributions were similar to Poisson distributions, which have been used in previous computational models of T6SS but have not yet been supported experimentally [15, 25].

Taken together, these results suggest that the subcellular T6SS dynamics in *V. fischeri*, including baseline activation level in liquid, rate of surface activation, and number of sheaths per cell, exhibits strain-specific variations. To systematically understand the effects of these strain-specific variations on the outcomes of intraspecific competition, we turn to mathematical and computational modeling.

### A biochemical model of T6SS

The biochemical model of T6SS consists of stochastic processes in two stages: (1) activation, and (2) structure assembly and deployment. This type of multi-stage model based on stochastic processes is an established approach for modeling transcription and translation [32, 33, 34, 35], and eukaryotic organelle synthesis [36].

#### Activation stage

We define activation as the cell being in a state ready to assemble T6SS structures. We denote the state of T6SS activation with *G*, such that *G*^−^ and *G*^+^ indicate the inactive and the active states, respectively. A cell in the inactive state *G*^−^ can be switched on at a constant rate λ_+_,

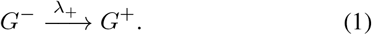

Here, we have abstracted into a single stochastic process the myriad processes that induce a T6SS activated state in a cell, e.g., sensing environmental signals which, depending on the system, could include copper or iron ions, temperature, or viscosity [37, 38, 28, 39], and translation and accumulation of T6SS proteins [40, 30, 41].

Based on experimental measurements on unprimed cultures (Fig. 2B), we introduce two additional parameters: *p*_0_, which accounts for the initial proportion of activated cells, and *τ*_+_, which accounts for the initial waiting period. Consider an infinitely large population of simple cell-like reactors that do not grow or divide, each of which independently undergoes activation as described by Eq. (1). At *t* = 0, each reactor has a probability *p*_0_ to be T6SS activated, and an inactive reactor waits *τ*_+_ before commencing stochastic switching. The activated percentage of the population *P* (*t*) at time *t* ≥ 0 is

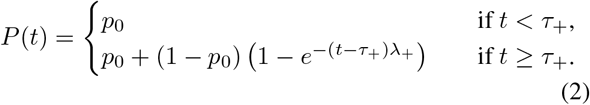

For both ES401 and FQ-A002, the measurements of activated population percentage at 0.5 h and 1 h in Fig. 2B are averaged to obtain their respective po values, and all measurements are used in nonlinear least square regressions to estimate their respective λ_+_ and τ_+_ (all values reported in Fig. 2 caption).

#### Structure assembly and deployment stage

The second stage in the T6SS biochemical model describes T6SS structure assembly and deployment. Previous studies in T6SS^+^ strains of *Pseudomonas aeruginosa, Vibrio cholerae*, and *Acinetobacter baylyi* have shown that the effects of T6SS structural proteins on T6SS assemblies can be categorized in two ways: the number of T6SS assemblies is driven by the abundance of proteins such as the spike protein VgrG and effector proteins, which are present in low copy numbers, whereas the length of the sheath in a functional structure mainly depends on the abundance of multimeric proteins such as Hcp and VipAB [40, 30, 41]. To incorporate these observations in our model, we correlate T6SS assembly with the appearance of a low abundance structural protein, which we assume is synthesized at a constant rate, λ_*s*_. We further assumed that, being in the activated state, all other component proteins are sufficiently abundant, thus do not limit the assembly. In terms of deployment, we simply consider that each T6SS structure can be independently fired at a constant rate, λ_*f*_. Since each functional T6SS structure has a sheath apparatus, henceforth, the term sheath is used interchangeably with T6SS structure. Thus, we have the following stochastic processes for the number of sheaths *N*,

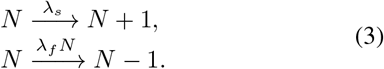

The T6SS biochemical model (Eqs. (1) & (3)) predicts that in a T6SS^+^ population with low initial T6SS activation, the average sheath number increases over time. The probability mass density function of number of sheaths per cell tends toward a steady state at long time. The rate of approach determined by λ_+_ and λ_*f*_ (SI Appendix Section 3, Fig. S5). The steady state average number of sheaths is 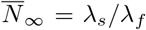, which implies that, at steady state, λ_*s*_ is the only parameter that controls the rate at which T6SS weapons can be produced and mobilized. We estimate 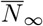 for ES401 and FQ-A002 by their respective experimental averages at 6 h (Fig. 2C).

In Eq. (1) we focus only on the activation process, and in Eqs. (3) we have neglected T6SS degradation processes independent of deployment. However, T6SS deactivation and degradation could be added if required. Although the T6SS biochemical model and its parameterization are informed by experimental results in lethal *V. fischeri* encoding T6SS2, it can be adapted to other T6SS regulatory systems.

### Integrated agent-based model (IABM)

T6SS activity at the subcellular level directly affects the ability of T6SS^+^ populations to kill, and thus influences the spatial structures on the length scale of the microbial colony [42, 15, 25]. To investigate the multi-scale interplay between subcellular T6SS dynamics, cellular growth, and intercellular interactions, we integrate the T6SS biochemical model into a custom ABM, resulting in an integrated ABM (IABM). In addition to imposing T6SS-dependent interaction rules among dueling cells as in some existing ABMs [42, 15, 25], each cell in our IABM to undergoes internal stochastic reactions (Eqs. (1) & (3)) governing the supply and usage of the T6SS arsenal. We treat each cell as a spherocylinder (a cylinder with hemispherical ends) growing in a monolayer on a viscous substrate. Cell growth is modeled as elongation along the cylinder axis according to the adder model [43], while the radius of the cell is kept constant. As the cells grow and come into contact with one another, the mechanical interactions cause them to move and rotate.

Each cell maintains the internal state variables *G* and *N*, T6SS activation state and sheath number, respectively, and carries out internal stochastic reactions (Eqs. (1) & (3)) at each time step in the simulation. If a cell fires a sheath, the target is randomly selected among the neighboring cells in contact. However, it is also possible for the cell to miss the neighbors and fire into the intercellular milieu. If the target is a clonemate, it survives; if the target is a nonclonal competitor cell, it ceases cellular function but participates in the mechanistic interactions for a time *τ*_lys_ until it completes lysis and disintegrates. At division, the daughter cells inherit the mother cell’s T6SS activation state G, and the mother cell randomly distributes its sheaths with equal probability to the two daughters. The internal T6SS reactions are also coupled to a cell’s physiology via growth. In maintaining a T6SS arsenal, we assume that the energetic cost of T6SS protein expression and assembly is dominant over the cost of firing structures, and maintaining the T6SS genes in the genome. A T6SS active cell has a penalized growth rate 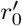 that decreases linearly with T6SS production rate λ_*s*_, i.e.,

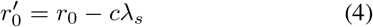

where *r*_0_ is the base growth rate of the strain if it did not produce T6SS and *c* is the cost coefficient. More details of the IABM are provided in SI Appendix Section 4.

### Model predictions and comparison with experimental data

#### Slow activation rate limits T6SS effectiveness

Results in the previous section (Fig. 2) indicate that the relatively slower surface activation rate in FQ-A002 is an important factor that led to the observation that wildtype FQ-A002 is outcompeted by wildtype ES401 under unprimed condition (Fig. 1A–D). To test whether our model captures this experimental observation, we simulate competitions between ES401 and FQ-A002 and between their *vasA* mutants under unprimed and primed conditions, as in the experiments presented in Fig. 1A–D. We create different computational strains to represent ES401, FQ-A002, and their *vasA*^−^ counterparts, based on previously estimated parameters with several adjustments. T6SS parameters for simulated wildtype ES401 are (*p*_0_, λ_+_, λ_*s*_, λ_*f*_, τ_lys_)=(10%, 0.6 h^−1^, 21 h^−1^, 7 h^−1^,0.5 h), and parameters for simulated wildtype FQ-A002 are (*p*_0_, λ_+_, λ_*s*_, λ_*f*_, τ_lys_)=(5%, 0.25 h^−1^, 21 h^−1^, 5.25 h^−1^, 0.5 h). To simulate a *vasA*^−^ mutant strain, we set λ_*f*_ = 0 in corresponding wildtype strain so that the mutant cells cannot attack using T6SS but still pay a growth penalty for expressing T6SS proteins. More details on IABM parameterization in **Materials and Methods** and SI Appendix Tables S1–S4.

Using a square periodic domain, we simulate the interior of the colony in a wildtype ES401 vs. FQ-A002 coincubation in both unprimed and primed conditions for an equivalence of 24 h. These simulations support our hypothesis, qualitatively matching the spatial characteristics of both unprimed and primed biological assays (Fig. 1A–B; SI Appendix Fig. S1BC; SI movies S1, S2). Simulated co-incubations of ES401 *vasA*^−^ vs. FQ-A002 *vasA*^−^ also capture the well-mixed spatial structure as observed in the corresponding biological assays (Fig. 1C; SI Appendix Fig. S1D).

We also hypothesize that the same factor (slow surface activation) leads to FQ-A002 being less effective than ES401 in eliminating nonlethal targets. To test this hypothesis, we simulate wildtype vs. *vasA*^−^ pairs as in the experiments presented in Fig. 1E, using two types of boundary conditions that represent either a confined space or a range expansion. As shown in Fig. 3A, ES401 *vasA*^−^ is able to grow and maintain its population for a longer time, compared to FQ-A002 *vasA*^−^, when coincubated with the wildtype lethal competitor. At later times 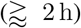, ES401 *vasA*^−^ and FQ-A002 *vasA*^−^ exhibit similar trends in the change in population size over time, differing by a multiplicative factor. Thus we confirm that wildtype FQ-A002, with slower activation rate, allows its nonlethal target, ES401 *vasA*^−^, to survive for longer and in larger numbers, qualitatively capturing the results in Fig. 1E.

**Figure 3:**
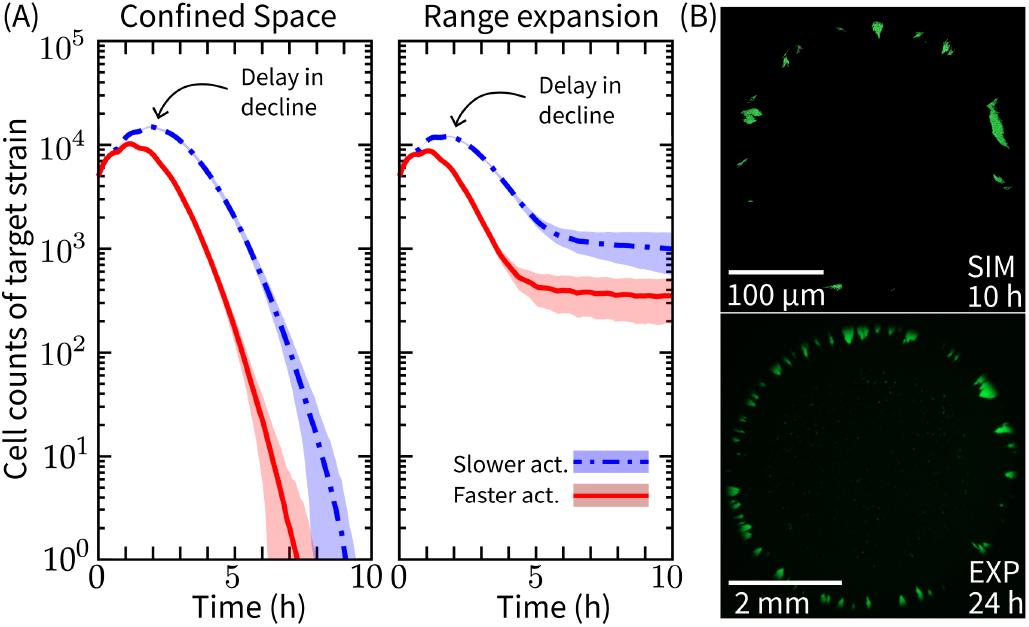
Target strains survive longer when competing against a slow activating lethal strains in a range expansion. (A) The cell counts of a target strain over time in when it is competed against various types of lethal strains, in a confined space (left) and a range expansion (right); results are averaged over 50 and 20 independent simulations, respectively. Colors correspond to the activation rate of the lethal strain in the coincubation simulations, and are consistent in both plots: blue = lethal strain activates slower with λ_+_ = 0.25 h^−1^, red = lethal strain activates faster with λ_+_ = 0.6 h^−1^. The shaded region on each curve show ±1 standard deviation. (B) Top: a representative simulation image of the slow activating (λ_+_ = 0.25 h^−1^) lethal (hidden) vs. target (green) coincubation in range expansion. Scale bar is 100 μm. See SI movie S3 and SI Appendix Table S3 for more simulation details. Bottom: a fluorescence microscopy image of the ES114 target strain following a coincubation with the ES401 inhibitor strain; scale bar is 2 mm. Unprimed strains were mixed at a 1:1 ratio and coincubated for 24 h on LBS agar plates.

#### Spatial environment of competition affects target survival

Bacteria compete in many different arenas, including environments where cells have space to expand their range as competition occurs, and those where competition is limited to a confined space, e.g. host colonization sites. In simulating wildtype vs. *vasA*^−^ competitions, we observe that the spatial environment plays a significant role in determining the survival of the target strain. In a confined spatial geometry, the target population becomes completely eliminated by the wildtype strain in 5 h–9 h (Fig. 3A, left). However, if the coincubation is allowed to grow in a range expansion, we find that a small population of target cells remain even after several hours of coincubation (Fig. 3A, right). This is because, even though the lethal cells can eliminate all target cells in the interior of the colony, a small number of target cells can persist in microcolonies at the edge of the coincubation spot (Fig. 3B, top). Only target cells at the boundary of these microcolonies come into contact with lethal cells, allowing the target cells bordered by clonemates to grow into open territory and reproduce. The survival of target cells at the edge of a colony has also been consistently observed in the laboratory assays of lethal vs. target coincubation (Fig. 3B, bottom).

### Cost determines the competitive fitness of T6SS production strategies

Variations in growth rates among bacterial strains can have significant impacts on competitive outcomes. While natural variations are common across bacterial strains, in this section, we focus our attention on isolating the effect of a growth penalty due to T6SS activity. We consider intraspecific competitions among two fully activated lethal strains with identical cost coefficient *c* and base growth rate *r*_0_, but different T6SS production rate λ_*s*_. We may expect a trade-off between the cost of T6SS production and its benefit in eliminating competitors.

We perform a parameter sweep using the IABM to understand the competitive fitness landscape in the context of the T6SS growth cost. We simulate a lethal resident strain with a fixed high T6SS production rate of λ_*s*,res_ = 20 h^−1^, and a competitor strain with varying T6SS production rate λ_*s*,comp_ ∈ [0, λ_*s*,res_]. To measure lethality of the competitor strain, we use a dimensionless parameter *β* = λ_*s*,comp_/(λ_*s*,res_+λ_*s*,comp_). For the range of λ_*s*,comp_ that we consider, *β* ∈ [0,0.5]. We also rescale the cost coefficient as *ĉ* = *c*/*c_max_* ∈[0, 1], where *c*_max_ = *r*_0_/λ_*s*,res_. To configure the competitor strain, we uniformly sample the parameter space of {*β*, *ĉ*|0<*β*<0.5,0<*ĉ*<1}. We assess the competitive outcome by *ϕ*=(*N*_res_ – *N*_comp_)/(*N*_res_+*N*_comp_), where *N*_res_ and *N*_comp_ are final cell counts of the resident and competitor strain, respectively. *ϕ* = 1,0, −1 indicate resident strain dominance, coexistence, and competitor strain dominance, respectively.

In Fig. 4A, we show competitive outcome *ϕ* as a function of *β* and *ĉ*. In region along the diagonal of parameter space of {*β*, *ĉ*}, the two competing strains can have different λ_*s*_ values but still coexist because they strike a similar balance in the growth vs. T6SS production trade-off. Above this diagonal region, the competitor strain (low producer) dominates, and below this diagonal, the resident strain (high producer) dominates. When *β* ≈ 0.5, both strains coexist at any value of *ĉ* due to comparable T6SS production levels.

**Figure 4:**
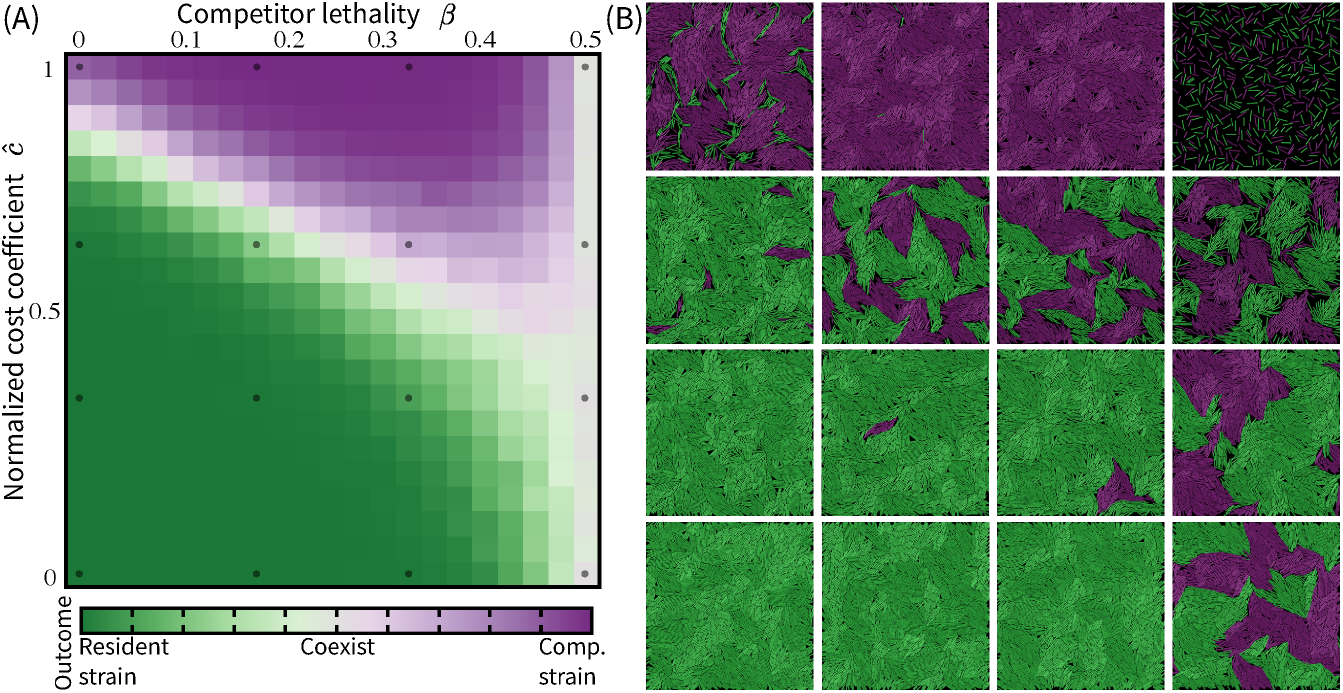
Competitive outcomes and surviving mechanisms are affected by the cost of T6SS production. (A) Phase space of competitive outcome in mutually lethal competitions as a function of two dimensionless parameters, *ĉ* and *β*, where *ĉ* is the normalized cost coefficient and *β* characterizes the lethality of the competitor strain. All simulations are run for an equivalence of 10 h. Final cell counts *N*_res_ and *N*_comp_, for the resident and competitor strain, respectively, are collected and averaged over 50 independent trials for each parameter combination {*β*, *ĉ*}. The competitive outcome is determined by *ϕ* = (*N*_res_ – *N*_comp_)/(*N*_res_ + *N*_comp_), with *ϕ* = 1, 0, −1 indicating resident strain dominance, coexistence, and competitor strain dominance, respectively. (B) Representative simulation images from across the parameter space. For other simulation parameters see SI Appendix Table S4.

Comparing simulations from across the parameter space (Fig. 4B), we find that there are various mechanisms through which one strain can dominate or coexistence can be achieved. When the cost of producing T6SS is low, i.e., *ĉ* < 1, the resident strain dominates by having a higher T6SS production, suppressing the faster growing competitor that maintains a smaller T6SS arsenal (Fig. 4B, lower left). With cost being low, when two strains coexist, cells grow and interact in T6SS-dependent manners to form spatially separated microcolonies, similar to what we observed in primed coincubations of wildtype ES401 vs. wildtype FQ-A002 (Fig. 1B; Fig. 4B, center to lower right). When the cost is high, i.e., *ĉ* ≈ 1, the competitor strain dominates by outgrowing the resident strain, which pays a heavy price for producing many T6SS structures (Fig. 4B, top left to center). With cost being high, cell growth can become extremely slow in high producers; thus two competition strains coexist because the initial population barely grow to establish intercellular contacts (Fig. 4B, top right).

## Discussion

T6SS is a highly dynamic system that is regulated and utilized by a wide range of microbes in diverse habitats, including animal hosts. Leveraging experimental data of surface-dependent T6SS activity in *V. fischeri*, our T6SS biochemical model captures the two most salient features of T6SS dynamics at the subcellular level that impact competition outcomes: the temporal variation of the activation period and the regulation of number of T6SS structures within the cell. The model is general and parsimonious, so that it can be applied to T6SSs in other microorganisms. More importantly, it can serve as the basis on which more complex T6SS biochemistry can be modeled and investigated.

By integrating the T6SS biochemical model into a custom ABM, we can simulate tens of thousands of cells growing, undergoing internal T6SS dynamics, and interacting mechanistically and in T6SS-dependent ways. Using the IABM, we demonstrate that intraspecific variations in activation, structure assembly, and deployment give rise to drastically different competitive outcomes between two T6SS^+^ strains under different competition conditions. We also apply the IABM to study the trade-off between the biological cost of T6SS structure production and its competitive benefits.

The variations in target survival when competition occurs in a confined space compared to a range expansion scenario provides insight into how competition might occur within a host. The simulation result suggests that if multiple strains enter a confined host colonization site, the target population may be quickly eliminated since it cannot avoid contact with the lethal strain. These results underscore the importance of considering the geometry of the competitive environment, which in addition to T6SS chemical dynamics at the subcellular level, can influence competitive outcomes.

## Materials and methods

### Media and growth conditions

*V. fischeri* strains were grown in LBS medium at 24 °C, and antibiotics were added to media for *V. fischeri* selection as described previously [44]. For selection in *V. fischeri* cultures, chloramphenicol, kanamycin, and erythromycin were added to LBS medium at final concentrations of 2 μg ml^−1^, 100 μg ml^−1^, and 5 μg ml^−1^, respectively.

### Coincubation assays

*V. fischeri* strains containing the indicated plasmid or chromosomal markers were grown overnight on LBS agar plates supplemented with the appropriate antibiotic at 24 °C. For each biological replicate, overnight cultures were started from a single colony and grown overnight in LBS supplemented with the appropriate antibiotic. Prior to the start of coincubation assays, cultures were either subcultured once more into liquid LBS for 6 h (unprimed treatments) or spotted onto an LBS agar plate for 6 h (primed treatments) as indicated. For each coincubation, strains were normalized to an OD_600_ of 1.0, mixed at a 1:1 ratio, and 5 μl of the mixture was spotted on LBS agar plates and incubated at 24 °C. At the indicated time points, coincubation spots were either imaged using fluorescence microscopy or total CFU counts were quantified using serial dilutions.

### Fluorescence microscopy

Fluorescence microscopy images of coincubation spots were imaged with a trinocular zoom stereo microscope equipped with a Nightsea fluorescence adapter kit for green and red fluorescence detection. Images were taken using an OMAX 14MP camera with OMAX ToupView camera control software. Single-cell images of VipA_2-GFP sheaths were taken either on an upright Olympus BX51 microscope outfitted with a Hammamatsu C8484-03G01 camera and a 100X/1.3 Oil Ph3 objective lens, or on an inverted Nikon Ti2 microscope outfitted with a Hammamatsu ORCA Fusion sCMOS camera and a CFI plan apo lambda 100X oil objective lens. Brightness and contrast adjustments were made uniformly across all images in a given experiment and color changes were made by adjusting the LUT value to either “green” or “magenta” in FIJI.

### Parametrization of simulated ES401 and FQ-A002 in the IABM

We use τ_+_ = 1 h for both simulated ES401 and FQ-A002 strains since the activated percentage in either biological ES401 or FQ-A002 population starts to increase after 1 h, and the difference in τ_+_ estimates for the two strains is below our experimental time resolution (Fig. 2B). The rate of surface activation in a biological strain depends on the experimental conditions. The particular experiments used to quantify this parameter made use of a sealed Petri dish for imaging (Fig. 2B), which likely limited oxygen supply to the bacteria. As the experimental setups in Fig. 1 and Fig. 2AC did not have this caveat, we increase the estimated activation rates of both ES401 and FQ-A002 (Fig. 2) by five fold while maintaining the ratio between the two. We base the value of λ_*s*_ ≈ 21 h^−1^ in our computational strains on publicly available microscopy video data that visualize sheath assembly and firing in *V. cholerae* [31]. The strain specific firing rates are determined by 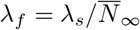. Since we have estimated 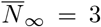 for ES401 and 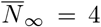 for FQ-A002 (Fig. 2C), the firing rates for the two strains are λ_*f*_ = 7 h^−1^, λ_*f*_ = 5.25 h^−1^, respectively. For τ_lys_, we use an intermediate value, 0.5 h, which is between the extremum values in a previous study that investigates the effect of lysis speed on T6SS effectiveness [15]. Furthermore, we let λ_s_ and τ_lys_ be identical in both simulated ES401 and FQ-A002 strains, since the two biological strains exhibit similar killing rates in coincubation with nonlethal targets (Fig. 1E). To simulate unprimed cells, after seeding the initial population at the start of the simulation, we randomly set a cell to be activated with probability *p*_0_, and let the inactive cells undergo stochastic switching as described in Eq. (1). In contrast, to simulate primed cells, we set every cell in the initial population to be activated at the start. The base growth rate *r*_0_ and T6SS growth penalty c are adjusted so that the penalized growth rate 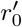 of the lethal strains are similar to those of biological strains ES401 and FQ-A002 [7]. For a summary of all IABM parameters, see SI Appendix Table S1; for parameters used in simulations in Figs. 1, 3, & 4, see SI Appendix Tables S2, S3, & S4, respectively.

## Acknowledgement

Y.L.L. acknowledges support for the Department of Energy Computational Science Graduate Fellowship, the NSF-Simons Center for Mathematical and Statistical Analysis of Biology at Harvard, award number #1764269 and the Harvard Quantitative Biology Initiative. S.S. was supported by the Department of Defense through the National Defense Science & Engineering Graduate Fellowship Program. A.N.S. was supported by NIGMS grant R35 GM137886. C.H.R. was partially supported by the Applied Mathematics Program of the U.S. DOE Office of Science Advanced Scientific Computing Research under contract number DE-AC02-05CH11231.

## Author contributions

Y.L.L, S.N.S., A.N.S., E.A.K., & C.H.R. conceptualized and designed the research, S.N.S. & A.N.S. designed experiments, S.N.S. conducted experiments and analyzed data, C.H.R. developed the initial agent-based model (ABM), Y.L.L, E.A.K., & C.H.R. adapted the ABM for T6SS systems, Y.L.L. developed the subcellular T6SS model, simulated the integrated ABM, and analyzed data, Y.L.L, S.N.S., A.N.S., E.A.K., & C.H.R. wrote the manuscript.

## Code availability

The simulation codes, BacSim-T6SS, are available on GitHub at https://github.com/ylunalin/BacSim-T6SS

## References

[1] Charles Darwin and Kebler Leonard. On the origin of species by means of natural selection, or, The preservation of favoured races in the struggle for life. J. Murray, London, 1859.

[2] Elisa T. Granato, Thomas A. Meiller-Legrand, and Kevin R. Foster. The evolution and ecology of bacterial warfare. Current Biology, 29(11):R521–R537, 2019.

[3] Lay-Sun Ma, Abderrahman Hechani, Jer-Sheng Lin, Alain Filloux, and Lai Erh-Min. *Agrobacterium tumefaciens* deploys a superfamily of type vi secretion dnase effectors as weapons for interbacterial competition in planta. Cell host & microbe, 16(1):94–104, July 2014.

[4] Thibault G. Sana, Nicolas Flaugnatti, Kyler A. Lugo, Lilian H. Lam, Amanda Jacobson, Virginie Baylot, Eric Durand, Laure Journet, Eric Cascales, and Denise M. Monack. *Salmonella* typhimurium utilizes a t6ss-mediated antibacterial weapon to establish in the host gut. Proceedings of the National Academy of Sciences, 113(34):E5044–E5051, 2016.

[5] Adrian J. Verster, Benjamin D. Ross, Matthew C. Radey, Yiqiao Bao, Andrew L. Goodman, Joseph D. Mougous, and Elhanan Borenstein. The Landscape of Type VI Secretion across Human Gut Microbiomes Reveals Its Role in Community Composition. Cell Host & Microbe, 22(3):411–419.e4, September 2017.

[6] Yang Fu, Brian T. Ho, and John J. Mekalanos. Tracking *Vibrio cholerae* cell-cell interactions during infection reveals bacterial population dynamics within intestinal micro environments. Cell host & microbe, 23(2):274–281.e2, February 2018.

[7] Lauren Speare, Andrew G. Cecere, Kirsten R. Guckes, Stephanie Smith, Michael S. Wollenberg, Mark J. Mandel, Tim Miyashiro, and Alecia N. Septer. Bacterial symbionts use a type VI secretion system to eliminate competitors in their natural host. Proceedings of the National Academy of Sciences, 115(36):E8528–E8537, September 2018.

[8] Jordan Vacheron, Maria Péchy-Tarr, Silvia Brochet, Clara Margot Heiman, Marina Stojiljkovic, Monika Maurhofer, and Christoph Keel. T6ss contributes to gut microbiome invasion and killing of an herbivorous pest insect by plant-beneficial *Pseudomonas protegens*. ISME journal, 13(5):1318–1329, May 2019.

[9] Andrew I. Perault, Courtney E. Chandler, David A. Rasko, Robert K. Ernst, Matthew C. Wolfgang, and Peggy A. Cotter. Host adaptation predisposes *Pseudomonas aeruginosa* to type vi secretion system-mediated predation by the *Burkholderia cepacia* complex. Cell Host & Microbe, 28(4):534–547.e3, August 2020.

[10] Petr G. Leiman, Marek Basler, Udupi A. Ramagopal, Jeffrey B. Bonanno, J. Michael Sauder, Stefan Pukatzki, Stephen K. Burley, Steven C. Almo, and John J. Mekalanos. Type vi secretion apparatus and phage tail-associated protein complexes share a common evolutionary origin. Proceedings of the National Academy of Sciences, 106(11):4154–4159, 2009.

[11] David Veesler and Christian Cambillau. A Common Evolutionary Origin for Tailed-Bacteriophage Functional Modules and Bacterial Machineries. Microbiology and Molecular Biology Reviews, 75(3):423–433, September 2011.

[12] Abdelrahim Zoued, Yannick R. Brunet, Eric Durand, Marie-Stéphanie Aschtgen, Laureen Logger, Badred-dine Douzi, Laure Journet, Christian Cambillau, and Eric Cascales. Architecture and assembly of the type vi secretion system. Biochimica et Biophysica Acta (BBA) - Molecular Cell Research, 1843(8):1664–1673, 2014.

[13] Alistair B. Russel, S. Brook Peterson, and Joseph D. Mougous. Type vi secretion system effectors: poisons with a purpose. Nature Reviews Microbiology, 12:137–148, February 2014.

[14] Amy J. Gerc, Andreas Diepold, Katharina Trunk, Michael Porter, Colin Rickman, Judith P. Armitage, Nicola R. Stanley-Wall, and Sarah J. Coulthurst. Visualization of the Serratia Type VI Secretion System Reveals Unprovoked Attacks and Dynamic Assembly. Cell Reports, 12(12):2131–2142, September 2015.

[15] William P. J. Smith, Andrea Vettiger, Julius Winter, Till Ryser, Laurie E. Comstock, Marek Basler, and Kevin R. Foster. The evolution of the type VI secretion system as a disintegration weapon. PLOS Biology, 18(5):e3000720, May 2020.

[16] Luke P. Allsopp, Patricia Bernal, Laura M. Nolan, and Alain Filloux. Causalities of war: The connection between type vi secretion system and microbiota. Cellular Microbiology, 22(3):e13153, 2020.

[17] Giuseppina Mariano, Katharina Trunk, David J. Williams, Laura Monlezun, Henrik Strahl, Samantha J. Pitt, and Sarah J. Coulthurst. A family of Type VI secretion system effector proteins that form ion-selective pores. Nature Communications, 10(1):5484, December 2019.

[18] Luke McNally, Eryn Bernardy, Jacob Thomas, Arben Kalziqi, Jennifer Pentz, Sam P. Brown, Brian K. Hammer, Peter J. Yunker, and William C. Ratcliff. Killing by Type VI secretion drives genetic phase separation and correlates with increased cooperation. Nature Communications, 8:14371, February 2017.

[19] D. Volfson, S. Cookson, J. Hasty, and L. S. Tsimring. Biomechanical ordering of dense cell populations. Proceedings of the National Academy of Sciences, 105(40):15346–15351, October 2008.

[20] Pushpita Ghosh, Jagannath Mondal, Eshel Ben-Jacob, and Herbert Levine. Mechanically-driven phase separation in a growing bacterial colony. Proceedings of the National Academy of Sciences, 112(17):E2166–E2173, April 2015.

[21] Pahala Gedara Jayathilake, Prashant Gupta, Bowen Li, Curtis Madsen, Oluwole Oyebamiji, Rebeca GonzÃ!lez-Cabaleiro, Steve Rushton, Ben Bridgens, David Swailes, Ben Allen, A. Stephen McGough, Paolo Zuliani, Irina Dana Ofiteru, Darren Wilkinson, Jinju Chen, and Tom Curtis. A mechanistic Individual-based Model of microbial communities. PLOS ONE, 12(8):e0181965, August 2017.

[22] Rafael D. Acemel, Fernando Govantes, and Alejandro Cuetos. Computer simulation study of early bacterial biofilm development. Scientific Reports, 8(1), December 2018.

[23] David Bruce Borenstein, Peter Ringel, Marek Basler, and Ned S. Wingreen. Established Microbial Colonies Can Survive Type VI Secretion Assault. PLOS Computational Biology, 11(10):e1004520, October 2015.

[24] Jared L. Wilmoth, Peter W. Doak, Andrea Timm, Michelle Halsted, John D. Anderson, Marta Ginovart, Clara Prats, Xavier Portell, Scott T. Retterer, and Miguel Fuentes-Cabrera. A Microfluidics and Agent-Based Modeling Framework for Investigating Spatial Organization in Bacterial Colonies: The Case of Pseudomonas Aeruginosa and H1-Type VI Secretion Interactions. Frontiers in Microbiology, 9:33, February 2018.

[25] William P. J. Smith, Maj Brodmann, Daniel Unterweger, Yohan Davit, Laurie E. Comstock, Marek Basler, and Kevin R. Foster. The evolution of tit-for-tat in bacteria via the type VI secretion system. Nature Communications, 11(1):5395, December 2020.

[26] Clotilde Bongrand and Edward G. Ruby. Achieving a multi-strain symbiosis: strain behavior and infection dynamics. ISME journal, 13(3):698–706, March 2019.

[27] Alecia N. Septer. The *Vibrio*-Squid Symbiosis as a Model for Studying Interbacterial Competition. mSystems, 4(3):e00108–19, June 2019.

[28] Lauren Speare, Stephanie Smith, Fernanda Salvato, Manuel Kleiner, and Alecia N. Septer. Environmental Viscosity Modulates Interbacterial Killing during Habitat Transition. mBio, 11(1):e03060–19, February 2020.

[29] Christina C. Saak, Martha A. Zepeda-Rivera, and Karine A. Gibbs. A single point mutation in a TssB/VipA homolog disrupts sheath formation in the type VI secretion system of *Proteus mirabilis*. PLOS ONE, 12(9):e0184797, September 2017.

[30] Lin Lin, Emmanuelle Lezan, Alexander Schmidt, and Marek Basler. Abundance of bacterial Type VI secretion system components measured by targeted proteomics. Nature Communications, 10(1):2584, December 2019.

[31] M. Basler, M. Pilhofer, G. P. Henderson, G. J. Jensen, and J. J. Mekalanos. Type VI secretion requires a dynamic contractile phage tail-like structure. Nature, 483(7388):182–186, March 2012.

[32] David R. Rigney. Stochastic model of constitutive protein levels in growing and dividing bacterial cells. Journal of Theoretical Biology, 76(4):453–480, February 1979.

[33] Johan Paulsson. Models of stochastic gene expression. Physics of Life Reviews, 2(2):157–175, June 2005.

[34] Ryszard Rudnicki and Andrzej Tomski. On a stochastic gene expression with pre-mRNA, mRNA and protein contribution. Journal of Theoretical Biology, 387:54–67, December 2015.

[35] Mukund Thattai. Universal Poisson Statistics of mRNAs with Complex Decay Pathways. Biophysical Journal, 110(2):301–305, January 2016.

[36] Shankar Mukherji and Erin K O’Shea. Mechanisms of organelle biogenesis govern stochastic fluctuations in organelle abundance. eLife, 3:e02678, June 2014.

[37] C. S. Bernard, Y. R. Brunet, E. Gueguen, and E. Cascales. Nooks and Crannies in Type VI Secretion Regulation. Journal of Bacteriology, 192(15):3850–3860, August 2010.

[38] Julie M. Silverman, Yannick R. Brunet, Eric Cascales, and Joseph D. Mougous. Structure and Regulation of the Type VI Secretion System. Annual Review of Microbiology, 66(1):453–472, October 2012.

[39] Martina Lazzaro, Mario F. Feldman, and Eleonora Garcìa Véscovi. A Transcriptional Regulatory Mechanism Finely Tunes the Firing of Type VI Secretion System in Response to Bacterial Enemies. mBio, 8(4):e00559–17, /mbio/8/4/e00559-17.atom, September 2017.

[40] Andrea Vettiger and Marek Basler. Type VI Secretion System Substrates Are Transferred and Reused among Sister Cells. Cell, 167(1):99–110.e12, September 2016.

[41] Chih-Feng Wu, Yun-Wei Lien, Devanand Bondage, Jer-Sheng Lin, Martin Pilhofer, Yu-Ling Shih, Jeff H Chang, and Erh-Min Lai. Effector loading onto the VgrG carrier activates type vi secretion system assembly. EMBO reports, 21(1), January 2020.

[42] Peter David Ringel, Di Hu, and Marek Basler. The Role of Type VI Secretion System Effectors in Target Cell Lysis and Subsequent Horizontal Gene Transfer. Cell Reports, 21(13):3927–3940, December 2017.

[43] Ariel Amir. Cell size regulation in bacteria. Phys. Rev. Lett., 112:208102, May 2014.

[44] Eric V. Stabb, Karl A. Reich, and Edward G. Ruby. *Vibrio fischeri* genes *hvnA* and *hvnB* encode secreted nad+-glycohydrolases. Journal of Bacteriology, 183(1):309–317, 2001.

