## Supplemental appendix for "A subcellular biochemical model for T6SS dynamics reveals winning competitive strategies"

#### 1 Additional experimental and simulation data for main text Fig. 1

##### 1.1 Competition outcomes vary due to strain-specific variations in T6SS surface activation response

In the main text, we introduce two treatments of inocula to prepare them for competition assays (main text Fig. 1). In the unprimed treatment, inocula are raised in liquid culture, mixed, and spotted directly on to the agar surface for competitive coincubation. In contrast, inocula can also be primed prior to mixing and competing by a clonal incubation period on an agar surface. Priming on a viscous surface activates and enhances T6SS structure formation and T6SS-dependent killing in *V. fischeri* [1]. We also perform competition assays between *vasA*$^-$ strains under either treatment. Since *vasA* gene disruption blocks the expression of a T6SS structural component, mutants with this disruption cannot make functional T6SS even though they still express majority of the T6SS-associated proteins. These assays serve as controls, showing that factors besides T6SS-dependent interactions do not give rise to the spatial organization and population levels observed in wildtype vs. wildtype competitions under the corresponding conditions. Whether the coincubations are under primed or unprimed conditions, competition outcomes between *vasA*$^-$ strains of ES401 and FQ-A002 are visually similar.

In Fig. 1 of the main text, we report results of *vasA* mutants of ES401 and FQ-A002 competing under the unprimed treatment. In Fig. S1A, we show an example microscopy image of a control assay under primed treatment. We also show four additional simulation samples for unprimed wildtype vs. wildtype competition (Fig. S1B), primed wildtype vs. wildtype competition (Fig. S1C), and *vasA*$^-$ vs. *vasA*$^-$ competitions (Fig. S1D). In *vasA*$^-$ vs. *vasA*$^-$ IABM simulations, because T6SS is not engaged in intercellular interactions, we do not distinguish between primed or unprimed conditions since the algorithmic details are identical in both scenarios.

Finally, we include two simulation movies:

**Movie S1** Unprimed wildtype ES401 vs. wildtype FQ-A002 competition. **Movie S2** Primed wildtype ES401 vs. wildtype FQ-A002 competition.

##### 1.2 Lethal strains ES401 and FQ-A002 have similar growth rates in lethal vs. target coincubations

In the main text, we describe experiments where we coincubated lethal vs. target pairs of (1) wildtype ES401 vs. FQ-A002 *vasA*$^-$ , and (2) wildtype FQ-A002 vs. ES401 *vasA*$^-$ (Fig. 1E in the main text). There, we observed that ES401 *vasA*$^-$ was able to maintain its population level during $1\,\mathrm{h} - 2\,\mathrm{h}$, whereas the CFU counts of FQ-A002 *vasA*$^-$ continued to decline during the same time window. This is consistent our hypothesis that strain-specific variations in T6SS surface activation response affects population dynamics of the interacting bacterial strains. In particular, this result supports our claim that FQ-A002, having a slower surface activation response, allows its targets to survive longer and in larger numbers. To rule out different growth rates being the reason for the observed difference in target decline in main text Fig. 1E, we present data of the colony forming units (CFU) of the lethal strains in each coincubation in Fig. S2, showing that the growth rates of both lethal strains are similar in these coincubation assays.

#### 2 Spatial environment of competition affects target survival

In the main text we show that nonlethal targets can survive T6SS attacks from lethal competitors in a range expansion, where target cells are protected by clonemates and can grow into territories free of lethal cells. In addition to the snapshot we provide in Fig. 3 in the main text, we provide a simulation movie that demonstrates this phenomenon of target cells surviving on the colony edges.

**Movie S3** Target cells survive T6SS attacks on the edges of a range expansion.

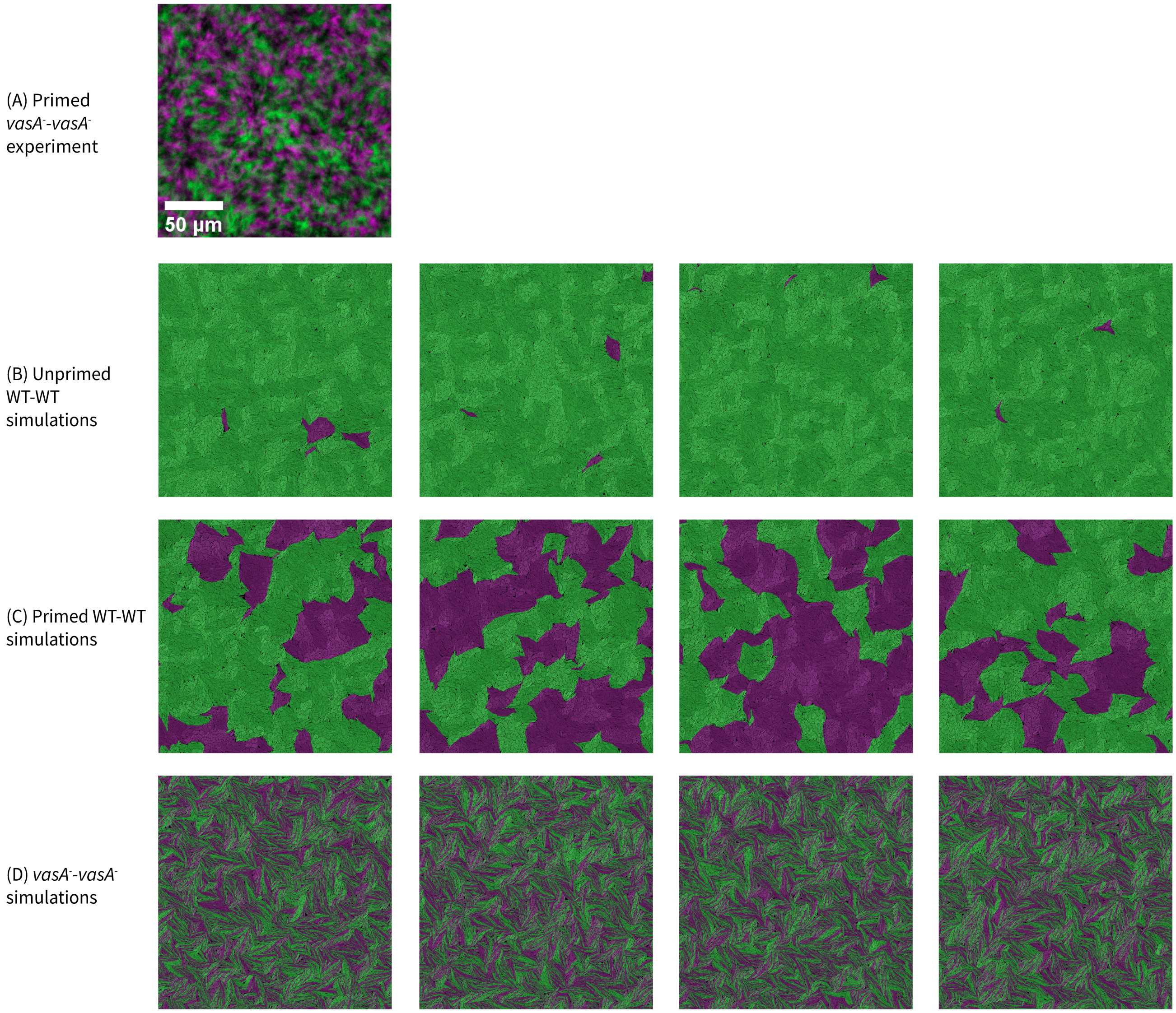


Figure S1: **Additional experimental and simulation data on wildtype vs. wildtype, *vasA*$^-$ vs. *vasA*$^-$ coincubations under unprimed and primed conditions.** (A) Example microscopy image of primed *vasA*$^-$ vs. *vasA*$^-$ competition, which serves as a control assay in the experiments described in Fig. 1A–D in the main text. Experimental protocol see corresponding sections in the main text. $T = 24\,\mathrm{h}$ images of four independent simulations of unprimed wildtype vs. wildtype simulations (B), primed wildtype vs. wildtype simulations(C), and *vasA*$^-$ vs. *vasA*$^-$ competitions (D).

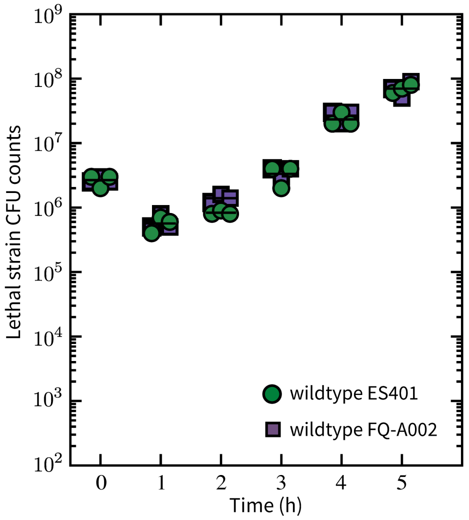


Figure S2: **Lethal wildtype strains ES401 (green) and FQ-A002 (purple) exhibit similar growth rate in lethal vs. target coincubations.** Wildtype ES401 is coincubated with FQ-A002 *vasA*$^-$ , wildtype FQ-A002 is coincubated with ES401 *vasA*$^-$ . Both pairs are grown under the unprimed condition. Other experimental protocols are reported in Fig. 1E and the **Materials and Methods** section in the main text.

### 3 T6SS biochemical model predicts shifting of the sheath number distribution

Consider the two-stage T6SS biochemical model,

$$\begin{aligned} G^- &\xrightarrow{\lambda_+} G^+, \\ N &\xrightarrow{\lambda_s} N{+}1, \\ N &\xrightarrow{\lambda_f N} N{-}1, \end{aligned} \tag{S1}$$

where $N$ is the number of T6SS structures (sheaths), and the binary state variable $G$ represents the cells being in T6SS active ($G^+$) or inactive ($G^-$) state. T6SS structures are produced at a constant rate $\lambda_s$ and fired at a rate proportional to the number of structures, $\lambda_f N$.

Consider a cell in the activated state $G^+$, in a small time interval $(t, t+\delta)$, the probability of producing a sheath is $\lambda_s(\delta + o(\delta))$. The probability of a sheath being fired is $N\lambda_f(\delta+o(\delta))$. The probability of not producing any sheath is $1-\lambda_s(\delta+o(\delta))$, that of not using any sheath is $1 - N\lambda_f(\delta + o(\delta))$. Therefore, the probability of neither producing nor using any structure is

$$(1 - \lambda_s(\delta + o(\delta)))(1 - N\lambda_f(\delta + o(\delta))) \approx 1 - \delta(\lambda_s + N\lambda_f) \tag{S2}$$

where we have assumed the probability of having two or more reactions within $\delta$ is small, i.e. $\lim_{\delta\to 0} \frac{o(\delta)}{\delta} = 0$. Denote the probability of having $N = n$ structures at time $t$ as $P_n(t)$. Then

$$\begin{aligned} P_n(t+\delta) &= P_{n-1}(t)(\lambda_s(\delta + o(\delta))) + P_{n+1}(t)((n+1)\lambda_f(\delta + o(\delta))) \\ &\quad + P_n(t)(1 - \lambda_s(\delta + o(\delta)) - n\lambda_f(\delta + o(\delta))) \\ &\approx \lambda_s\delta P_{n-1}(t) + (n+1)\lambda_f\delta P_{n+1}(t) + P_n(t)(1 - \lambda_s\delta - n\lambda_f\delta). \end{aligned} \tag{S3}$$

We approximate time derivative of $P_n(t)$ by taking the limit of $\delta \to 0$,

$$\begin{aligned} \frac{\partial P_n(t)}{\partial t} &= \lim_{\delta\to 0} \frac{P_n(t+\delta) - P_n(t)}{\delta} \\ &= -(\lambda_s + n\lambda_f)P_n(t) + \lambda_s P_{n-1}(t) + (n+1)\lambda_f P_{n+1}(t). \end{aligned} \tag{S4}$$

This is the master equation satified by the probability mass function of the number of sheaths, for $n \geq 0$. Next we will derive the governing equation for the probability generation function of this probability distribution, and solve for the generating functions for $\lambda_f = 0$ and $\lambda_f > 0$. The generating function for a probability distribution is

$$G(s,t) = \sum_{n=-\infty}^{\infty} s^n P_n(t). \tag{S5}$$

We will use the following expressions in our derivations:

$$\sum_{n=-\infty}^{\infty} s^n n P_n(t) = s\frac{\partial G(s,t)}{\partial s}, \tag{S6}$$

$$G(1,t) = \sum P_n(t) = 1, \tag{S7}$$

$$\langle n \rangle = \left.\frac{\partial G(s,t)}{\partial s}\right|_{s=1} \equiv G'(1,t), \tag{S8}$$

$$\langle n(n-1) \rangle = \left.\frac{\partial^2 G(s,t)}{\partial s^2}\right|_{s=1} \equiv G''(1,t), \tag{S9}$$

$$\langle n^2 \rangle = \left.\frac{\partial}{\partial s}\left(s\frac{\partial G(s,t)}{\partial s}\right)\right|_{s=1}. \tag{S10}$$

To proceed, we make use of a step operator, which is a linear operator commonly used in solving the master equations such as Eq. (S4). Details of techniques for handling master equations such as the generating function and the step operator can be found in standard textbooks for stochastic processes such as Ref. [2]. The linear step operator $E^l[\cdot]$ where $l \in \mathbb{Z}$ can be applied to a function $f(n)$, where the support $n$ can run over the real numbers. The linear operator can be defined as

$$\begin{aligned} E^l[f(n)] &= f(n+l), \\ (E^l - 1)[f(n)] &= f(n). \end{aligned} \tag{S11}$$

Step operations have the property

$$\sum_{n=-\infty}^{\infty} s^n (E^k - 1)[f(n)] = (s^{-k} - 1) \sum_{n=-\infty}^{\infty} s^n f(n). \tag{S12}$$

Rewriting the time derivative of the probability distribution in Eq. (S4) with the linear step operator we introduced in Eqs. (S11), we have

$$\frac{\partial P_n(t)}{\partial t} = (E-1)[n\lambda_f P_n(t)] + (E^{-1} - 1)[\lambda_s P_n(t)]. \tag{S13}$$

Taking the partial time derivative of Eq. (S5) and using Eq. (S13), we have

$$\begin{aligned} \frac{\partial G(s,t)}{\partial t} &= \sum_{n=-\infty}^{\infty} s^n \left\{(E-1)[n\lambda_f P_n(t)] + (E^{-1}-1)[\lambda_s P_n(t)]\right\} \\ &= (s^{-1} - 1) \sum_{n=-\infty}^{\infty} s^n (n\lambda_f P_n(t)) + (s-1) \sum_{n=-\infty}^{\infty} s^n (\lambda_s P_n(t)) \\ &= \lambda_f (1-s)\frac{\partial G(s,t)}{\partial s} + \lambda_s (s-1) G(s,t). \end{aligned} \tag{S14}$$

In the second line, we have used the property of jump functions, Eq. (S12), and in the third line, we have used Eq. (S6) for the first term, and the definition of $G(s,t)$ in Eq. (S5) for the second term. The last equation above, Eq. (S14), is a first-order partial differential equation (PDE) for $G(s,t)$, which can be solved by the method of characteristics, with an appropriate initial condition

$$G(s,0) = G_0(s). \tag{S15}$$

For $\lambda_f = 0$, Eq. (S14) reduces to an ordinary different equation. Solving the initial value problem with the initial condition in Eq. (S15) yields

$$G(s,t) = e^{\lambda_s (s-1)t} G_0(s). \tag{S16}$$

Differentiating $G(s,t)$ with respect to $s$ and evaluating the derivatives appropriately (Eqs. (S8) & (S10)), we find that the mean and the variance of the number of T6SS structures are

$$\langle N(t)\rangle = \left.\frac{\partial G(s,t)}{\partial s}\right|_{s=1} = \lambda_s t + \langle N(0)\rangle, \tag{S17}$$

$$\sigma_N^2(t) = \frac{\partial}{\partial s}\left(s\frac{\partial G(s,t)}{\partial s}\right)\Bigg|_{s=1} - \langle N(t)\rangle^2 = \lambda_s t + \sigma_N^2(0), \tag{S18}$$

respectively, where $N(0)\rangle = G_0'(1)$ and

$$\sigma_N^2(0) = \langle N(0)^2\rangle - \langle N(0)\rangle^2 = G_0''(1) + G_0'(1) - (G_0'(1))^2. \tag{S19}$$

In the expression of initial variance, $\sigma_N^2(0)$, we have used identities in Eqs. (S8) & (S9). For $\lambda_f > 0$, Eq. (S14) has a solution

$$G(s,t) = \exp\left\{\tfrac{\lambda_s}{\lambda_f}(s-1)(1-e^{-\lambda_f t})\right\} G_0((s-1)e^{-\lambda_f t}+1). \tag{S20}$$

Differentiating Eq. (S20) shows that the mean is

$$\langle N(t)\rangle = \left.\frac{\partial G(s,t)}{\partial s}\right|_{s=1} = \tfrac{\lambda_s}{\lambda_f}(1-e^{-\lambda_f t}) + e^{-\lambda_f t}\langle N(0)\rangle \tag{S21}$$

and the variance is

$$\begin{aligned}\sigma_N^2(t) &= \frac{\partial}{\partial s}\left(s\frac{\partial G(s,t)}{\partial s}\right)\Bigg|_{s=1} - \langle N(t)\rangle^2 \\ &= \left(\tfrac{\lambda_s}{\lambda_f} + e^{-\lambda_f t}\langle N(0)\rangle\right)(1-e^{-\lambda_f t}) + e^{-2\lambda_f t}\sigma_N^2(0).\end{aligned} \tag{S22}$$

Taken together, Eqs. (S20), (S21) & (S22) indicate that in a system with $\lambda_f > 0$, the effect of the initial condition is damped out exponentially at a rate of $\lambda_f$. The probability mass distribution of sheaths is a Poisson distribution with time-dependent mean. In the long time limit, the steady state distribution has the mean $\bar{N}_{\text{st}} = \lambda_s/\lambda_f$.

### 4 Combine T6SS biochemical model with a 2D agent-based model

In this section, we introduce the agent-based model and discuss the integration of the T6SS biochemical model (Eqs. (S1)). The agent-based model (ABM) is designed for microbial growth on two-dimensional (2D) surfaces such as agar plates, and it takes into account cell growth, division, and cell–cell and cell–substrate interactions. Under fast growth conditions, the model of cell growth and division implemented in the ABM reduces to the so-called adder model [3]. However, in surface growth, factors such as pressure induced by crowding, nutrient depletion, and waste accumulation limit cell growth, leading to an increase in doubling time, or stopping growth altogether. To simulate contact-dependent T6SS interactions on a surface, it is necessary to model surface growth. We will discuss the mechanism we use to constrain cell growth in this scenario.

#### 4.1 Single cell ideal growth

We represent a cell as a spherocylinder with two hemispherical caps of radius $R$, and a body length $l_{\text{cyl}}$. The total cell length is $l = l_{\text{cyl}} + 2R$. We represent the growth of such a cell by elongation along its cylindrical axis, and use $l$ to represent cell size in lieu of cell volume. In a population of cells with an average growth rate $r_0$, the average doubling time is $\tau_c = \ln(2)r_0^{-1}$. Let $l$ and $l_b$ denote an individual's current cell length and length at birth, respectively. Cells are shown to grow exponentially, therefore, $l(t) = l_b e^{tr_0}$. Let $l_0$ be the population average cell length at birth, and $l_d$ be the length at time of division. A unified model of single cell size regulation has been proposed [3]. The cell size at division is

$$l_d = 2^{1+\frac{\eta_t}{\tau_c}} l_b^{1-a} l_0^a, \tag{S23}$$

where $\eta_t$ is a noise in interdivision time, and parameter $a$ gives rise to several common size regulation models. For example, in the absence of noise, choosing $a = 1$ results in a model where cells divide at a critical length, i.e. $l_d = 2l_0$. This is called the sizer model because it results in tight control of birth sizes around average size $l_0$. Choosing $a = 0$ leads to constant growth time, so that cells always double birth size, i.e. $l_d = 2l_b$. This is called the timer model because the growth time is tightly distributed around the mean. In the ABM, we choose $a = 0.5$ which results in the so-called adder model. In this growth model, cells add a constant volume regardless birth size. Therefore, cells that are larger than average at birth grow for a time shorter than average cell cycle, and vice versa. The adder model produces statistics that agree with experiments, in particular, coefficients of variance between size at birth and size at division, and distributions of cell size and interdivision time. For details, see the works of Amir and coworkers and references therein [3, 4, 5, 6].

In simulating cell cycle using Eq. (S23), we use a small Gaussian noise $\eta_t \sim \mathcal{N}(0, \sigma_t^2)$ to perturb the interdivision time. Division is considered symmetric between the daughter cells, however, a small Gaussian noise in length $\eta_l \sim \mathcal{N}(0, \sigma_l^2)$ is used to perturb the daughter cell sizes at birth. To illustrate this, consider a mother cell of length $l_d$, symmetrically dividing into two daughters. Daughter 1 and daughter 2 have cell length at birth

$$l_{b,1} = \tfrac{1}{2} l_d + \eta_l, \qquad l_{b,2} = l_d - l_{b,1}, \tag{S24}$$

respectively. After division, we perturb the orientations of daughter cells by a small Gaussian noise $\eta_\theta \sim \mathcal{N}(0, \sigma_\theta^2)$, to avoid artificial chaining. Values of the noise size parameters are summarized in Table S1.

#### 4.2 Mechanical interactions and growth restriction in the ABM

Within a population from the same species or strain, growth rates can vary from individual to individual due to idiosyncratic variations. Since we simulate cells growing on a two-dimensional surface, the most prominent factor that limits the growth is space availability. Therefore, we employ a pressure-based growth restriction factor to effectively stop cells from elongating when the space is depleted. To facilitate this discussion, we first describe mechanical interactions among cells, followed by the details of the growth restriction model.

##### 4.2.1 Interactions within a population

As cells grow and divide, they experience mechanical forces from other cells as they come into contact and push on one another. Cells also experience damping forces due to contact with the viscous substrate. Let us denote cells in the ABM by $b_i, b_j, b_k, \ldots$. We assume cells to be made of a linear elastic material with elastic modulus $E$, which has the dimension of force per area. When cells get pushed into each other either due to growth, or due to mechanical interactions with neighboring cells, the intercellular contact between a pair of cells $i$ and $j$ causes a small deformation $h$ in both cells. As shown in the schematic in Fig. S3(A), the region enclosed by dashed curves indicates the total deformation needed to accommodate the pairwise contact. The maximum width $H$ of this region, which can also be viewed as overlap between two perfectly rigid spherocylinders, can be computed by standard contact detection methods. The deformation of each cell $h$ is $h = \frac{1}{2}H$, since two contacting cells equally deform in the process. The strain in each cell can be obtained by scaling with a typical length $\frac{h}{R}$. Assuming that the contact area scales as $\sim R^2$,* the magnitude of the repulsive force exerted on cell $i$ due to cell $j$ scales as $|\mathbf{f}_{ij}| \sim ER^2 \frac{h}{R} = ERh$. The direction along which the repulsive forces are acting in this pair of cells can be determined from the geometry as illustrated in Fig. S3(A). We sum over all such forces on a cell and the viscous force it experiences to get the total force on this cell, $\mathbf{F}_i$.

The Newtonian dynamics of cell $i$ is described by the equations of motion,

$$\begin{aligned} \dot{\mathbf{x}}_i &= \mathbf{v}_i, \\ m_i \dot{\mathbf{v}}_i &= \sum_{j \in J} \mathbf{f}_{ij} - \gamma_l \mathbf{v}_i, \end{aligned} \tag{S25}$$

where $\mathbf{x}_i$ and $\mathbf{v}_i$ are the position and velocity of cell $i$, respectively. The cell mass is $m_i = \rho(\pi R^2 l_{i,\text{cyl}} + \frac{4}{3}\pi R^3)$. $\gamma_l$ is the viscous damping coefficient of the substrate, $\mathbf{f}_{ij}$ denotes the contact forces exerted by cell $j$ on $i$, where $j$ is in the set of cells that are in contact with $i$, denoted by $J$. The contact forces also induce a torque on the cells. The angular dynamics is described by

$$\begin{aligned} \dot{\theta}_i &= \omega_i, \\ I_i \dot{\omega}_i &= \sum_{j \in J} \tau_{ij} - \gamma_r \omega, \end{aligned} \tag{S26}$$

*Our assumption of constant contact area leads to a Hookean contact model, a commonly-used model in discrete-element method simulations of particles [7]. A more accurate description would assume that contact area depends on $h$, which can lead to, e.g., the Hertzian contact model [8]. However the Hertzian model is more computationally expensive to implement, and has a limited effect on the dynamics of moving particle assemblies when compared to experimental results [9]. We therefore make use of the simpler Hookean model.

where $\theta_i$ and $\omega_i$ are the orientation and angular velocity of cell $i$, respectively. $\gamma_r$ is the viscous damping coefficient for rotating cell bodies, $I_i$ is the moment of inertia of a spherocylinder rotating about its center, perpendicular to the cylindrical axis,

$$I_i = \frac{2}{5} m_{i,\text{sph}} R^2 + m_{i,\text{sph}} (l_{i,\text{cyl}}/2)^2 + \frac{1}{12}\left(3R^2 + l_{i,\text{cyl}}^2\right) m_{i,\text{cyl}}, \tag{S27}$$

where $m_{i,\text{sph}} = \frac{4}{3}\rho R^3$ is the sum of the masses of the two hemispherical caps, and $m_{i,\text{cyl}} = \rho\pi R^2 l_{i,\text{cyl}}$ is the mass of the cylinder. In the equation above, the first term is the moment of inertia of a sphere, or that of two hemispheres. The second term applies the parallel axis theorem to translate the two hemispheres to either end of the cell. The third term is the moment of inertia of a cylinder. Using dimensional analysis [10], $\gamma_r \sim A\gamma_l$, where $A$ has the dimension of area. Equations (S25) & (S26) make up the system of ODEs that describes motion of a cell, and we solve these equations for each cell using an explicit forward Euler scheme.

#### 4.3 Parameter estimation

The exact values of $\gamma_l$ and $\gamma_r$ are not important, as long as the values provide sufficient damping in the system. We make use of an estimation procedure that provides reasonable values of these two parameters. In our mechanistic model above, we have the following parameters: growth rate $r_0$, average cell size $l_0$, cell radius $R$, cell density $\rho$, linear and rotational viscous damping coefficients $\gamma_l, \gamma_r$, and elastic modulus $E$. Using Buckingham's $\Pi$ theorem, we can construct four dimensionless groups. Using $\rho, l_0, R$ we can construct an average mass $m_0 = \rho(\pi R^2(l_0 - 2R) + \frac{4}{3}\pi R^3)$. Suppose a cell with mass $m_0$ is pushed by one end of an exponentially growing cell. If the motion is linear, i.e. the cell only undergoes translation, it experiences a pushing force due to the growing neighboring cell $\sim r_0^2 l_0$. The viscous damping force is $\sim \gamma_l r_0 l_0/m_0$. Taking the quotient of the two, we have a dimensionless parameter $p_1 = \gamma_l/m_0 r_0$, which is the ratio between viscous damping force and the intercellular force due to growth of neighboring cells. We set $p_1 = 100$, i.e. $\gamma_l = 100 m_0 r_0$, to so that any motion in the system is sufficiently damped out. To set $\gamma_r$, we use the dimensionless group $p_2 = \gamma_r/(A_0\gamma_l) = 1$ where $A_0 = (l_0 - 2R)R + \pi R^2$ is the largest horizontal cross-section area of a cell of average cell size. We assume that this is the typical area in contact with the viscous substrate.

Another dimensionless parameter, $p_3 = ER^2/(\gamma_l r_0 l_0)$, represents the ratio between elastic repulsive forces and the damping forces. This parameter helps us determine the timestep restriction in the simulation. We require that the large contact forces among cells be sufficiently damped out by the viscous interactions. Therefore, the maximum timestep is $dt_{\text{max}} = m_0/(p_3\gamma_l)$. The last dimensionless parameter $p_4 = ER^2/(m_0 r_0^2 l_0)$ can be interpreted as the elastic responses to contact and pushing due to cell growth. Parameters $p_3$ and $p_4$ are important in addressing a common issue in ABMs of cellular growth, which is that cells may exhibit unrealistic overlap after some time of growth. In a model where spatial configuration in the monolayer of cells is important to resolve contact-dependent killing, preventing this problem becomes important. We need to choose elastic modulus $E$ such that $p_3$ is sufficiently large to keep cells separate on the viscous substrate, and that $p_4$ is sufficiently large to keep cells separate when they grow. We find that setting $p_3 = 10$ is sufficient to prevent overlap among cells in our simulations, i.e. $E = 10\gamma_l r_0 l_0/R^2$. These dimensionless parameters are summarized Table S1, and values of dimensional parameters $dt, \gamma_l, \gamma_r$, and $E$, which depend on the physiology of the cells we simulate, are computed at the start of each simulation.

##### 4.3.1 Mechanistic growth restriction

To further address the problem of unrealistic overlap among cells in late stages of growth, we find it is also necessary to impose a growth limit. Although restrictions in growth can be due to resource availability, waste accumulations, antibacterial chemicals, etc., the primary factor we focus on is the availability of space. Intercellular pressure can reach a high value in a crowded region of the bacterial colony, which can slow down and even stop growth altogether [11, 12, 13]. The contact model described above provides us a convenient way to impose a growth constraint based on crowdedness, and a similar mechanism to constrain growth has been used in other ABMs [14].

We assume that inhibition of growth is due mechanical forces exerted along the growth axis, but independent of forces orthogonal to the axis (Fig. S3(B)). Let $r$ denote the restricted growth rate, $r = \beta r_0$, where $\beta \leq 1$ is a multiplicative growth restriction factor. Therefore, the actual interdivision time is $\tau_c = (\log 2)r^{-1}$, independent of the small noise. The expression for $\beta$ is

$$\beta = \max\left(0, 1 - \frac{P_{\text{up}}\, P_{\text{low}}}{P_c(P_{\text{up}} + P_{\text{low}})}\right) \tag{S28}$$

where $P_{\text{up}}$ and $P_{\text{low}}$ are calculated from contact forces at the upper and lower ends of a cell, respectively, and the critical pressure, $P_c$, determines when a cell stops growing. This form of growth restriction resembles the empirical model for growth restriction imposed by nutrient availability developed by Monod [15], and has the property that it requires pressure experienced by both ends of the cell to act together to stop growth. To give some physical intuition, if one end of the cell is blocked by neighbors, but the other end of the cell is free from contact with any cell, the cell should be able to grow toward the free side without any growth impediment. In fact, in this scenario, Eq. (S28) evaluates to 1. Similar to the half saturation constant in the Monod nutrient restriction model, $P_c$ can be considered a half inhibition constant—if the pressures experienced by both ends are $P_c$, then the cell growth rate is halved. The pressure along the growth (cylindrical) axis is computed by taking the projection of the contact force along the cell axis, and normalizing it by the total cross-sectional area of the cell, $\pi R^2$. The critical pressure $P_c$ is currently an unknown parameter but it must be determined *a priori*. In practice, we choose a small but sufficient value, $P_c = \xi_c E$ with $\xi_c = 5 \times 10^{-3}$, to address the issue while avoiding over-constraining cellular growth. Note that in the two-dimensional system, pressure has the dimension of force per length.

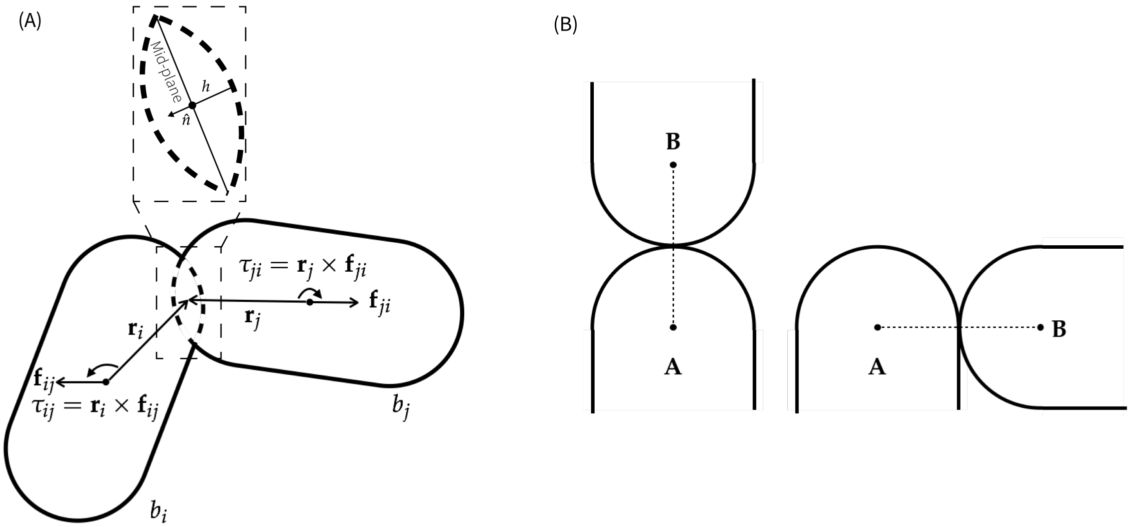


Figure S3: (A) Schematics of cell-cell interaction. The amount of deformation needed to accommodate two cells in contact can be considered as overlap between two spherocylinders (dashed region). The amount of deformation $h$ of each cell is half of the maximum width of the overlap region. $\hat{n}$ denotes the direction along which the repulsive forces are acting, it is perpendicular to the mid-plane of the overlap region. Also indicated in the schematics are contact forces, moment arms, and resulting torques, acting on each cell in contact. (B) Left: the contact force between cell A and cell B can potentially inhibit both cells, because pressure due to contact is along the growth axis of both A and B. If the other end of a cell is also blocked, the pressure from growth will build up and stop the cell from elongating. Right: the contact force between A and B would only contribute to cell B's growth inhibition. However, the equal but opposite contact force on cell A only applies a torque on the cell, and does not inhibit its growth.

#### 4.4 Integration of the T6SS biochemical model in the ABM

Here we expand on how we integrate the T6SS biochemical model into the cells' internal state in the agent-based model. The T6SS activation state $G$ and apparatus number $N$ are stored as internal state variables. As described in the main text, the activation probability is

$$P(t) = \begin{cases} p_0 & \text{if } t < \tau_0, \\ p_0 + (1 - p_0)\left(1 - e^{-(t-\tau_0)\lambda_+}\right) & \text{if } t \geq \tau_0, \end{cases} \tag{S29}$$

where the parameter $p_0$ accommodates the liquid culture activity level and $\tau_0$ takes into account a possible wait time (lag phase) before the activation process begins.

To incorporate these parameters in the simulation, each initial seed cell has a probability of $p_0$ to be active. If there is a waiting period $\tau_0 > 0$ (which can be strain-specific or idiosyncratic to each cell), then during a cell's waiting period $0 \leq t < \tau_0$

it does not engage in any biological processes. This means it does not, grow, divide, activate T6SS assembly, form T6SS structures, or fire any strucutre. After the waiting period, during each time step, each cell elongates its length according to

$$\frac{dl}{dt} = rl \tag{S30}$$

where $r$ is the actual growth rate specific to each cell, subject to constraint due to space limitation and cost of producing and using T6SS. We model the T6SS cost as a penalty on the base growth rate [14], so that a T6SS$^+$ cell that is active has growth rate

$$r_0' = r_0 - c\lambda_s \tag{S31}$$

where $c$ is the cost coefficient. Note that if a cell can produce and fire T6SS structures, but is in an inactive state, then the base growth rate is unaltered. Combined with mechanical growth restriction, the cell specific actual growth rate is $r = \beta r_0'$.

In addition, the following events can occur with appropriate probabilities according to Eqs. (S1):

1. an inactive cell becomes activated,
2. an active cell produces a new T6SS apparatus,
3. a cell that has any T6SS apparatuses can fire one.

We require that a cell must be activated before it can produce sheaths, and a cell can only fire if it has one or more sheaths. Therefore, in inactivated cells, event 1 is the only possible event with probability $\lambda_+ dt$. In activated cells, event 2 is always possible with probability $\lambda_s dt$, and event 3 is possible when the sheath number $N > 0$, with probability $\lambda_f N dt$.

Upon firing a T6SS apparatus, the attacking cell randomly selects one of its neighboring cells as the target, or fire into the environment without hitting any target. In our T6SS biochemical model the average number of sheaths per cell is regulated via a balance between production and usage. Translated to the ABM, we must simulate cells that can fire into the extracellular milieu even when it is isolated from any neighboring cells. In doing so, we avoid an excessive accumulation of sheaths within an isolated cell, which leads skewed sheath number distribution and increased firing frequency when the cell comes into contact with others later on.

To simulate this, we consider each neighbor cell and the extracellular milieu as a slot to fire into. The cell can fire T6SS attack into each slot with equal probability. For example, if the cell has two neighboring cells, then the T6SS is fired with equal probability $\frac{1}{3}$ into either of the neighbors or the extracellular milieu. After each cell has its turn in the T6SS-dependent interaction, we survey the entire simulation and determine if a cell is still alive. If the target is a sister cell of the attacker, it remains alive. Otherwise, it is marked as dead. A dead cell ceases growth, division, and other internal processes immediately, but still participates in the mechanical interactions in the simulation. An internal lysis timer starts counting from the moment a cell is marked as dead, and it is removed from the simulation after lysis is complete.

Division occurs at the end of a time step for any eligible cells. During division, a mother cell's sheaths are randomly distributed to the two daughter cells according to a binomial distribution. The probability of a daughter cell receiving $k$ out of $n$ sheaths, given its mother cell has $n$ sheaths, is

$$\mathbb{P}_d(k|n) = \binom{n}{k}\frac{1}{2^n}. \tag{S32}$$

The total probability of a newborn cell having $k$ sheaths needs to account for the probability of the mother cell having $n$ sheaths,

$$\mathbb{P}_d(k) = \sum_{n=0}^{\infty} \mathbb{P}_d(k|n)\mathbb{P}_m(n) \tag{S33}$$

$$= \sum_{n=0}^{\infty} \frac{n!}{k!(n-k)!}\frac{1}{2^n}\frac{\bar{N}^n e^{-\bar{N}}}{n!} \tag{S34}$$

$$= \frac{1}{k!}\left(\frac{\bar{N}}{2}\right)^k e^{-\bar{N}} \sum_{n=0}^{\infty} \frac{1}{(n-k)!}\left(\frac{N}{2}\right)^{n-k} \tag{S35}$$

$$= \frac{1}{k!}\left(\frac{\bar{N}}{2}\right)^k e^{-\bar{N}/2} \tag{S36}$$

where $\bar{N} = \lambda_s/\lambda_f$ is the steady-state mean of the Poisson distribution reflected in Eq. (S21). The subscripts $d$ and $m$ denote daughter and mother. Here we have assumed that the probability mass function of sheaths in mother cells has reached steady state. On a single cell level, we assume that the firing time scale is much faster than the cell cycle time, so that the exponentially decaying terms in Eqs. (S21) and (S22) can be treated as zero. However, division disrupts this equilibrium. The calculation above shows that the sheath number distribution of a population of cells immediately after division is also a Poisson distribution, but the division process halves the mean and variance. Following the discussion in Section 3, however, this initial distribution at birth equilibrates to the steady state Poisson distribution with the original mean and variance exponentially fast, with rates dependent on firing rate $\lambda_f$.

#### 4.5 Summary of IABM model units and parameters

The simulating unit length can be matched on to physical length by rescaling with average cell length of the bacteria. We estimate average cell size of our experimental strains from microscopy images, and report these values in Fig. S4. For simplicity, for all simulated strains we use $3\,\mu\text{m}$ for the average cell length at birth, $0.6\,\mu\text{m}$ for width, which stays constant during growth. In the simulation, the bacteria are spherocylinders with constant radius $1\,\text{L}$. Thus the simulation unit length can be converted to physical unit by $1\,\text{L} = 0.3\,\mu\text{m}$. We also let one unit simulation time be one hour in physical time, i.e. $1\,\text{T} = 1\,\text{h}$. Similarly, we can rescale the mass unit M in the IABM so that the simulation mass density $\rho$ has a value of unity, $1\,\text{M}/\text{L}^3$.

Previous experiments on which we base our experimental methods [16] estimate that the doubling time of a T6SS$^-$ strain of *V. fischeri*, ES114, is approximately $39\,\text{min} \pm 5.8\,\text{min}$. The doubling time of the T6SS$^+$ strain FQ-A002 is estimated to be approximately $43\,\text{min} \pm 10\,\text{min}$ under identical experimental conditions. We adopt these growth rate estimates, in particular, we use doubling time $39\,\text{min}$, i.e., growth rate $1.07\,\text{h}^{-1}$, as the base growth rate for all strains in the simulation. We further assume that the observed difference in doubling time can be attributed to diverting material and energy needed for growth to expressing T6SS-associated proteins, based on an estimate of $\lambda_s \approx 21\,\text{h}^{-1}$, we can use Eq. (S31) to estimate the default cost coefficient, $c \approx 0.005$.

We categorize and summarize all model parameters of the IABM in Table S1, and additionally provide parameters used in the simulations reported in Figs. 1, 3, & 4 in the main text, which may be be strain-specific and take on varying values (Tables S2, S3, & S4).

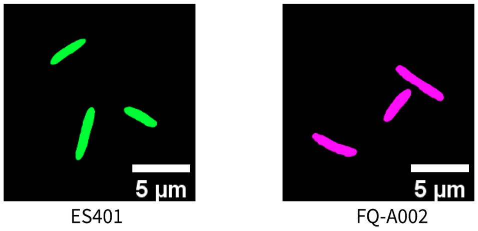


Figure S4: **Quantification of sizes of ES401 and FQ-A002 cells.** Overnight cultures of wildtype ES401 and FQ-A002 harboring the GFP expressing plasmid pVSV102 were grown clonally in unprimed conditions prior to spotting $2\,\mu\text{l}$ onto an agarose pad. Cells were imaged in the FITC channel within $10\,\text{min}$ of spotting; length and width measurements were calculated using NIS Elements software. For strain ES401, 533 cells were analyzed across 10 individual images. ES401 $3.2\,\mu\text{m} \pm 1.1\,\mu\text{m}$, width $0.64\,\mu\text{m} \pm 0.12\,\mu\text{m}$ (left). For strain FQ-002, 512 cells were analyzed across 10 individual images. FQ-A002 length $3.5\,\mu\text{m} \pm 1.3\,\mu\text{m}$, width $0.65\,\mu\text{m} \pm 0.11\,\mu\text{m}$ (right). Representative images are shown.

| Name | Symbol | Value | Unit |
|---|---|---|---|
| System parameters | | | |
| Domain size | $L$ | Varying | µm |
| Boundary condition | - | Periodic or free | - |
| Simulation time | $T$ | Varying | h |
| Initial cell number density | $\rho_{\text{num}}$ | 8.3 (default), varies | per $100\,\mu m^2$ |
| Ratio between damping forces and the intercellular forces due to growth | $p_1$ | 100 | dimensionless |
| Ratio between linear and rotational damping coefficients | $p_2$ | 1 | dimensionless |
| Ratio between elastic forces and damping forces | $p_3$ | 10 | dimensionless |
| Multiplier for critical pressure in growth rate regulation | $\xi_c$ | $5 \times 10^{-3}$ | dimensionless |
| Noise size in doubling time perturbation | $\sigma_t$ | 0.02 | h |
| Noise size in division length perturbation | $\sigma_l$ | 0.1 | µm |
| Noise size in post-division orientation perturbation | $\sigma_\theta$ | 0.01 | rad |
| Cell mass density | $\rho$ | $\sim 1$ | $g\,cm^{-3}$ |
| Cell physiology parameters | | | |
| Cell width | $R$ | 0.6 | µm |
| Average cell length at birth | $l_0$ | 3.0 | µm |
| Average base growth rate | $r_0$ | 1.07 | $h^{-1}$ |
| T6SS associated parameters | | | |
| Initial activation percentage (liquid culture activity level) | $p_0$ | Varies | dimensionless |
| Activation rate | $\lambda_+$ | Varies | $h^{-1}$ |
| Structure synthesis rate | $\lambda_s$ | Varies | $h^{-1}$ |
| Structure firing rate | $\lambda_f$ | Varies | $h^{-1}$ |
| Structure synthesis cost coefficient | $c$ | 0.005 (default), Varies | dimensionless |
| Lysis time | $\tau_{\text{lys}}$ | 0.5 (default), Varies | h |

Table S1: A list of simulation parameters in the custom integrated agent-based model.

| System parameters | | | | |
|---|---|---|---|---|
| Name | Symbol | Value | | Unit |
| Square domain size | $L$ | 400 | | µm |
| Boundary condition | - | Periodic | | - |
| Simulation time | $T$ | 24 | | h |
| Initial cell number density | $\rho_{\text{num}}$ | 8.3 | | per 100 µm$^2$ |
| **T6SS associated parameters** | | | | |
| Name | Symbol | ES401 | FQ-A002 | Unit |
| Initial activation percentage | $p_0$ | 10% | 5% | dimensionless |
| Activation rate | $\lambda_+$ | 0.6 | 0.25 | h$^{-1}$ |
| Structure synthesis rate | $\lambda_s$ | 21 | 21 | h$^{-1}$ |
| Structure firing rate | $\lambda_f$ | 7 | 5.25 | h$^{-1}$ |
| Structure synthesis cost coefficient | $c$ | 0.005 | | dimensionless |
| Lysis time | $\tau_{\text{lys}}$ | 0.5 | | h |

Table S2: Parameters used in the IABM simulations for Fig. 1 in the main text, and in Fig. S1B–D, for unprimed wildtype cells. To simulate primed cells, initial activation percentage $p_0$ is adjusted to $p_0 = 100\%$. To simulate *vasA*$^-$ cells, firing rate $\lambda_f$ is adjusted to $\lambda_f = 0$.

| System parameters | | | | |
|---|---|---|---|---|
| Name | Symbol | Range expansion | Confined spaces | Unit |
| Domain size | $L$ | 200 (initial spot radius) | 200 (square domain size) | µm |
| Boundary condition | - | Free | Periodic | - |
| Simulation time | $T$ | 10 | 10 | h |
| Initial cell number density | $\rho_{\text{num}}$ | 8.3 | 6.5 | per 100 µm$^2$ |
| **T6SS associated parameters** | | | | |
| Name | Symbol | Lethal strain | Target strain | Unit |
| Initial activation percentage | $p_0$ | 0% | 0% | dimensionless |
| Activation rate | $\lambda_+$ | 0.25 (slow), 0.6 (fast) | 0 | h$^{-1}$ |
| Structure synthesis rate | $\lambda_s$ | 21 | 0 | h$^{-1}$ |
| Structure firing rate | $\lambda_f$ | 7 | 0 | h$^{-1}$ |
| Structure synthesis cost coefficient | $c$ | 0.005 | 0 | dimensionless |
| Lysis time | $\tau_{\text{lys}}$ | 0.17 | 0 | h |

Table S3: Parameters used in the IABM simulations for Fig. 3 in the main text.

### 5 Activation curve and shifting sheath number distribution in integrated agent-based model simulations

As stated in Section 3, the sheath number distribution transitions from the initial distribution to a steady Poisson distribution by way of T6SS reactions in the absence of cell cycle. To verify that this transition in the distribution is also present in the agent-based model with internal T6SS dynamics, we simulate a clonal population of bacteria growing over time. We record the number of T6SS structures, or equivalently sheaths, per cell over time and the resulting histograms show that the transitory behavior in the sheath number distribution is indeed consistent with our analytical prediction. Simulation snapshots of a growing microcolony from a single bacterium, along with the activated population percentage and the time-dependent sheath number distributions, are shown in Fig. S5.

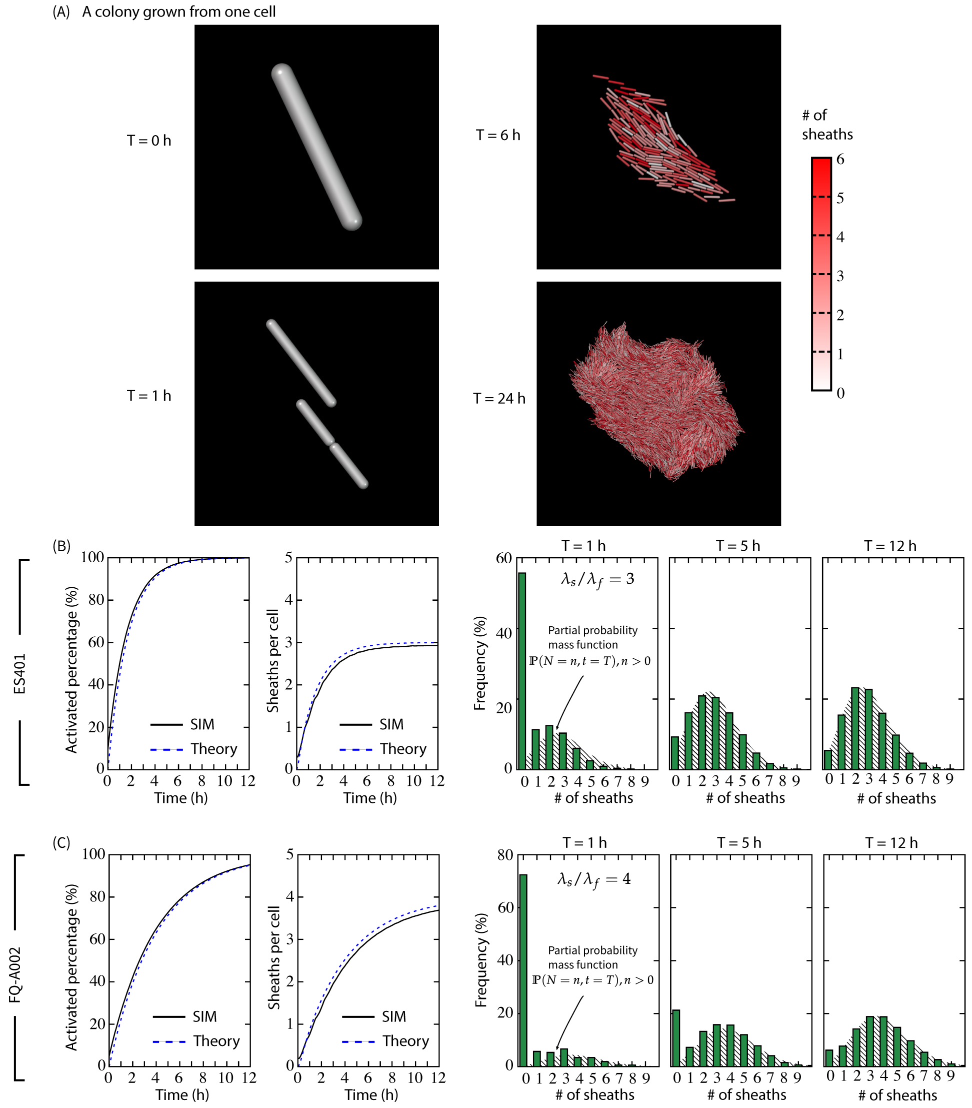


Figure S5: **Agent-based model confirms that sheath numbers distribution shifts over time in activating populations.** (A) Simulation snapshots of a growing microcolony from a single bacterium. Color indicates the number of sheaths a cell harbors. The simulations parameters are $(r_0, \lambda_+, \lambda_s, \lambda_f, p_0) = 0.924\,\mathrm{h}^{-1}, 1\,\mathrm{h}^{-1}, 45\,\mathrm{h}^{-1}, 15\,\mathrm{h}^{-1}, 0)$. Initially, cells have no sheaths. (B) Simulating a population of ES401 cells (in (C) FQ-A002 cells) growing and interacting mechanistically and in T6SS-dependent manner. We track the percentage of activated population and the number of sheaths per cell over time. Each simulation starts with 200 initial cells and grows for 12 h. Results from 200 simulations are averaged. The activation follows the theoretical prediction, $P(t) = 1 - \exp(-\lambda_+ t)$. The rate at which the average number of sheaths per cell tends to the steady state value is influenced by the activation processes. We compare simulation result with the expression $N(t) = \frac{\lambda_s}{\lambda_f}(1 - \exp(-\lambda_f t))P(t)$. We also predict the average number of sheaths per cell for ES401 is $\lambda_s/\lambda_f = 3$, ($\lambda_s/\lambda_f = 4$ for FQ-A002) and the steady state distribution is a Poisson distribution with that mean. Histograms of sheath numbers at $T = 1\,\mathrm{h}, 5\,\mathrm{h}, 12\,\mathrm{h}$ are plotted. At each time point, a fraction $P(t) \leq 100\%$ of the population is T6SS active. Since $\lambda_f \gg \lambda_+$, we can safely assume that within the sub-population that is T6SS active, the processes of producing and firing sheaths have reached steady state. Thus the probability mass function of having *no* sheath is $\mathbb{P}(N = 0, t) = (1 - P(t)) + P(t)e^{-\lambda_s/\lambda_f}$, and the probability of having $n > 0$ sheaths is $\mathbb{P}(N = n, t) = P(t)\frac{1}{n!}\left(\lambda_s/\lambda_f\right)^n e^{-\lambda_s/\lambda_f}$. The histograms in (B) and (C) are plotted against their respective partial probability mass function $\mathbb{P}(N = n, t = T), n > 0$ at each time $T$ when statistics are collected. Parameters $r_0, \lambda_+, \lambda_s, \lambda_f, p_0$ for ES401 (B) and FQ-A002 (C) are reported in Table S1.

| System parameters | | | | |
|---|---|---|---|---|
| Name | Symbol | Value | | Unit |
| Square domain size | $L$ | 78 | | μm |
| Boundary condition | - | Periodic | | - |
| Simulation time | $T$ | 10 | | h |
| Initial cell number density | $\rho_{\text{num}}$ | 8.3 | | per $100\,\mu\text{m}^2$ |
| **T6SS associated parameters** | | | | |
| Name | Symbol | Resident strain | Competitor strain | Unit |
| Initial activation percentage | $p_0$ | $100\%$ | | dimensionless |
| Structure synthesis rate | $\lambda_s$ | 20 | $[0, 20]$ | $\text{h}^{-1}$ |
| Structure firing rate | $\lambda_f$ | 20 | $[0, 20]$ | $\text{h}^{-1}$ |
| Structure synthesis cost coefficient | $c$ | $[0, 0.0533]$ | | dimensionless |
| Lysis time | $\tau_{\text{lys}}$ | 0.5 | | h |

Table S4: Parameters used in the IABM simulations for Fig. 4 in the main text.

### Supplmentary Information References


[1] Lauren Speare, Stephanie Smith, Fernanda Salvato, Manuel Kleiner, and Alecia N. Septer. Environmental Viscosity Modulates Interbacterial Killing during Habitat Transition. *mBio*, 11(1):e03060–19, February 2020.

[2] Toral Raúl. Introduction to master equations. In Pere Colet and Toral Raúl, editors, *Stochastic Numerical Methods*, pages 235–260. John Wiley & Sons, Ltd, 2014.

[3] Ariel Amir. Cell size regulation in bacteria. *Phys. Rev. Lett.*, 112:208102, May 2014.

[4] Po-Yi Ho and Ariel Amir. Simultaneous regulation of cell size and chromosome replication in bacteria. *Frontiers in Microbiology*, 6, July 2015.

[5] Felix Barber, Po-Yi Ho, Andrew W. Murray, and Ariel Amir. Details Matter: Noise and Model Structure Set the Relationship between Cell Size and Cell Cycle Timing. *Frontiers in Cell and Developmental Biology*, 5:92, November 2017.

[6] Po-Yi Ho, Jie Lin, and Ariel Amir. Modeling cell size regulation: From single-cell-level statistics to molecular mechanisms and population-level effects. *Annual Review of Biophysics*, 47(1):251–271, 2018. PMID: 29517919.

[7] Leonardo E. Silbert, Deniz Ertaş, Gary S. Grest, Thomas C. Halsey, Dov Levine, and Steven J. Plimpton. Granular flow down an inclined plane: Bagnold scaling and rheology. *Phys. Rev. E*, 64(5):051302, Oct 2001.

[8] K. L. Johnson. *Contact Mechanics*. Cambridge University Press, 1985.

[9] Chris H. Rycroft, Ashish V. Orpe, and Arshad Kudrolli. Physical test of a particle simulation model in a sheared granular system. *Phys. Rev. E*, 80:031305, Sep 2009.

[10] Grigory Isaakovich Barenblatt. *Scaling*. Cambridge University Press, 2003.

[11] B. I. Shraiman. Mechanical feedback as a possible regulator of tissue growth. *Proceedings of the National Academy of Sciences*, 102(9):3318–3323, March 2005.

[12] D. Volfson, S. Cookson, J. Hasty, and L. S. Tsimring. Biomechanical ordering of dense cell populations. *Proceedings of the National Academy of Sciences*, 105(40):15346–15351, October 2008.

[13] Da Yang, Anna D. Jennings, Evalynn Borrego, Scott T. Retterer, and Jaan Männik. Analysis of factors limiting bacterial growth in pdms mother machine devices. *Frontiers in Microbiology*, 9:871, 2018.

[14] William P. J. Smith, Andrea Vettiger, Julius Winter, Till Ryser, Laurie E. Comstock, Marek Basler, and Kevin R. Foster. The evolution of the type VI secretion system as a disintegration weapon. *PLOS Biology*, 18(5):e3000720, May 2020.

[15] Jacques Monod. The growth of bacterial cultures. *Annual Review of Microbiology*, 3(1):371–394, 1949.

[16] Lauren Speare, Andrew G. Cecere, Kirsten R. Guckes, Stephanie Smith, Michael S. Wollenberg, Mark J. Mandel, Tim Miyashiro, and Alecia N. Septer. Bacterial symbionts use a type VI secretion system to eliminate competitors in their natural host. *Proceedings of the National Academy of Sciences*, 115(36):E8528–E8537, September 2018.
